# The three-dimensional chromatin structure of the major human pancreatic cell types reveals lineage-specific regulatory architecture of T2D risk

**DOI:** 10.1101/2021.11.30.470653

**Authors:** Chun Su, Long Gao, Catherine L. May, James A. Pippin, Keith Boehm, Michelle Lee, Chengyang Liu, Matthew C. Pahl, Maria L. Golson, Ali Naji, the HPAP Consortium, Struan F.A. Grant, Andrew D. Wells, Klaus H. Kaestner

**Author notes:** Co-senior authors (Struan F.A. Grant, Andrew D. Wells, Klaus H. Kaestner).

## Abstract

Three-dimensional (3D) chromatin organization maps help to dissect cell type-specific gene regulatory programs. Furthermore, 3D chromatin maps have contributed to elucidating the pathogenesis of complex genetic diseases by connecting distal regulatory regions and genetic risk variants to their respective target genes. To understand the cell type-specific regulatory architecture of diabetes risk, we generated transcriptomic and 3D epigenomic profiles of human pancreatic acinar, alpha, and beta cells using single-cell RNA-seq, single-cell ATAC-seq, and high-resolution Hi-C of sorted cells. Comparisons of these profiles revealed differential A/B (open/closed) chromatin compartmentalization, chromatin looping, and transcriptional factor mediated control of cell type-specific gene regulatory programs. We identified a total of 4,750 putative causal-variant-target-gene pairs at 194 type 2 diabetes GWAS signals using pancreatic 3D chromatin maps. We found that the connections between candidate causal variants and their putative target effector genes are cell-type stratified and emphasize previously underappreciated roles for alpha and acinar cells in diabetes pathogenesis.

## Introduction

Type 2 diabetes (T2D) is a serious and costly disease that impacted an estimated 463 million people worldwide in 2019^1^. Disease onset is typically late (>40 years old), with incidence rising due to population aging and increasing rates of obesity. Of concern, more children and young adults are now developing type 2 diabetes^2^. Chronic complications include accelerated development of cardiovascular and microvascular disease; furthermore, diabetes is a major cause of blindness. While much of the increased prevalence of T2D can be attributed to the rising prevalence of obesity and a sedentary lifestyle, increasing evidence suggests that genetic risk factors underlie the pathogenesis of this common disease. As the genetic contribution to T2D is relatively weak and multifactorial when contrasted with that of type 1 diabetes (T1D)^3, 4^, it is more challenging to identify risk loci for T2D, earning the nickname of ‘geneticist’s nightmare’ prior to the advent of genome-wide association studies (GWAS).

GWAS have revolutionized the field of complex disease genetics and yielded compelling evidence for over 500 genetic loci associated with T2D^5^. The role of large consortia and international collaborations for achieving the scale and power necessary for larger GWAS analyses through the collection of material, combining genotype scores, and releasing data to the scientific community has greatly increased the number of risk loci uncovered for this disease. Unfortunately, GWAS only report genomic signals associated with a given trait and rarely the precise identity of the effector transcript and corresponding affected cell-types. As such, over the past 15 years GWAS has defined a period of signal discovery, rather than an era of gene target discovery.

Because gene expression in mammals is frequently controlled via long range interactions over large genomic distances, many regulatory elements do not control the nearest genes and can reside tens or hundreds of kilobases (kb) away. Therefore, attention is required when considering the biology underpinning an established GWAS signal, as frequently the gene nearest the lead SNP does not encode the effector transcript that impacts the phenotype. In addition, for complex diseases such as T2D, multiple organ systems contribute to the disease phenotype; in the case of diabetes, at a minimum this includes skeletal muscle, adipose tissue, liver, brain, and pancreas. Due to the predominant role of pancreatic islet regulatory sites for T2D risk^6^, three-dimensional chromatin maps in the whole human pancreatic islets and β cell cellular model EndoC-βH1 have been frequently used to annotateT2D risk signals to effector genes^7–9^. However, because islets consist of multiple cell types with distinct functions and different cell types can employ alternative *cis*-regulatory elements to control the same gene, it is necessary to further dissect variant-to-gene connections at pancreatic cell-type level to fully understand the molecular basic of T2D risk.

In this study, we combined 3D chromatin maps from the major pancreatic cell types with matched single cell transcriptome and chromatin accessibility data to investigate cell-type-specific gene regulation program in human pancreas. This approach also enabled mapping of more than 100 T2D GWAS signals to their likely effector transcript(s) and corresponding cell type(s) of action. Remarkably, in addition to the expected beta-cell acting variants, we also discovered variant-to-gene combinations that likely function in pancreatic alpha and acinar cells, suggesting an important contribution for these underappreciated cell types in T2D pathogenesis.

## Results

### Single-cell transcriptomic and chromatin accessibility profiles of the human pancreas

To explore the cell type-specific impact of chromatin architecture and accessibility on gene expression in the human pancreas, we performed high-resolution Hi-C (see Methods) on three distinct primary pancreatic cell subsets (acinar, alpha, and beta cells) FACS-sorted from multiple non-diabetic organ donors (**Supplemental Figure 1**). We then coupled this approach with scATAC-seq and scRNA-seq on whole pancreatic islets from three matched donors (**Figure 1A and Supplemental Table 1**) and performed integrative data analysis.

**Figure 1.**
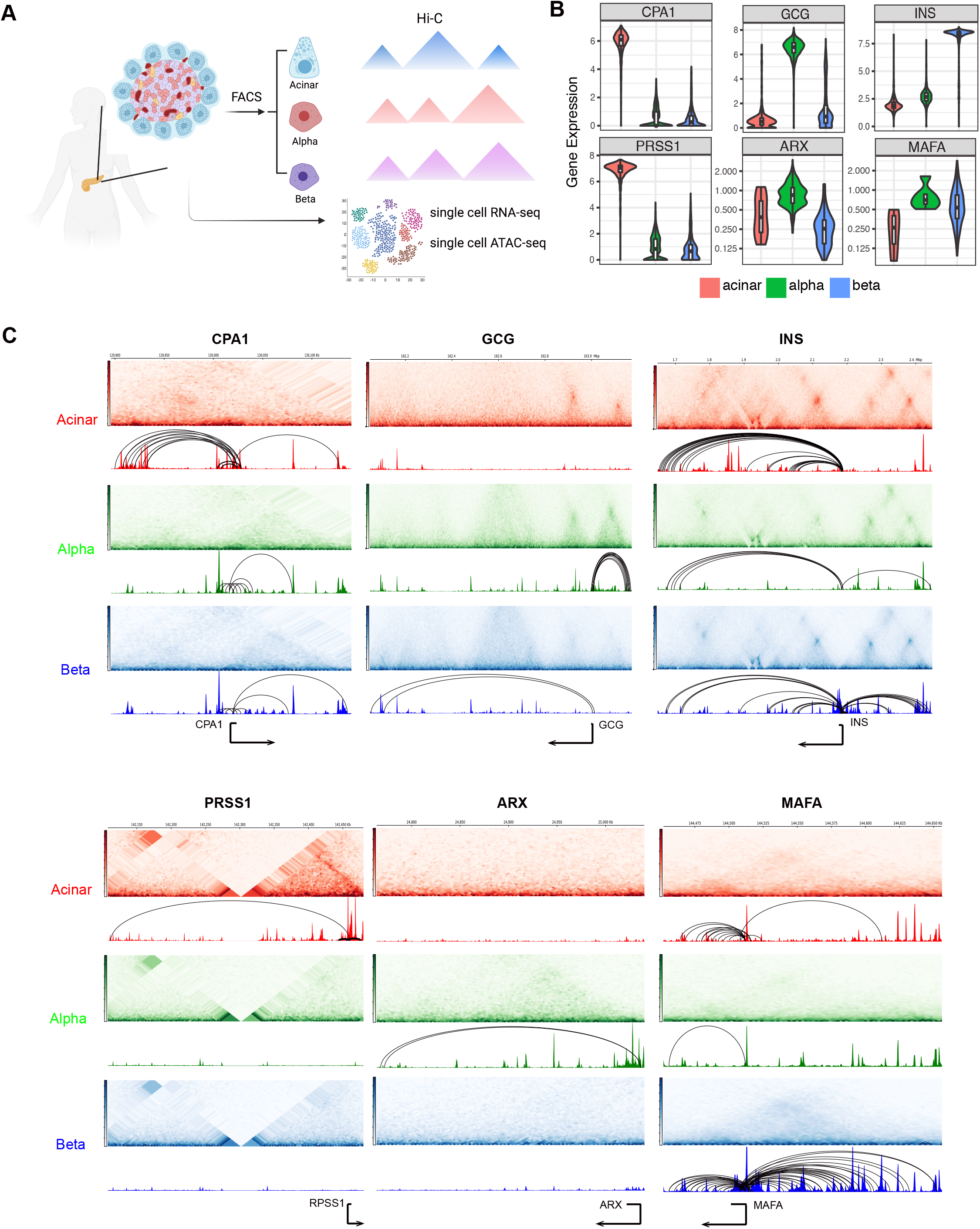
General validation of transcriptomic, chromatin accessibility and 3D architecture profiles in human pancreas cell subsets. **A**, Experimental design used in this study. Pancreatic islets obtained from 3 human donors were used for single cell experiments. Acinar, alpha and beta cells from 5 human donors were purified using FACS and followed by Arima Hi-C assays. See also **Supplemental Figure S1** and **Supplemental Tables S1**. B, Single-cell expression profiles for selected marker genes. Acinar: *CPA1* and *PRSS1*; alpha: *GCG* and *ARX*; beta: *INS* and *MAFA*. C, Aggregate single-cell chromatin accessibility, Hi-C contact frequency matrix and chromatin loops for selected marker genes.

To obtain high-quality, cell-type-specific transcriptomic profiles, we first clustered 5,288 quality-controlled single cells across the three donors from scRNA-seq into nine clusters (**Supplemental Figure 2A**) using conventional cell-type-specific markers (**Supplemental Figure 2B**). Donors were evenly represented across each of the identified cell types (**Supplemental Figure 2B**) with highly reproducible transcriptomic profiles (**Supplemental Figure 2C**). Acinar (19.8%), alpha (52.2%) and beta cells (16.1%) contributed to the three largest cell clusters from human pancreatic islets with distinctly expressed cell markers (**Figure 1B, Supplemental Figure 2B**). As previously observed^10^, endocrine cells (alpha, beta, delta, gamma and epsilon) formed a tight cluster to the exclusion of the other cell types, from which alpha and beta are closest to epsilon and delta cells, respectively, while acinar cells are closest to ductal cells in terms of transcriptomic profile (**Supplemental Figure 2D**). To confirm the purity of FACS-sorted cell populations, we performed bulk RNA-seq on sorted subsets of cells that were used for Hi-C library preparation and compared them to the single cell transcriptomic profile. On average, 93% of the sorted alpha gene program and 76% of the sorted beta cell program matched to the corresponding cell type cluster in scRNA-seq (**Supplemental Figure 2E**).

Next, we performed snATAC-seq on islet cells from the same set of donors and clustered accessible chromatin profiles from 12,473 high-quality single cells into nine cell type groups, similar to that observed with scRNA-seq (**Supplemental Figure 3A**). The promoter regions of marker genes showed high chromatin accessibility in the corresponding cell types, and each cell type was represented by all three donors (**Supplemental Figure 3B).** We further compared aggregated reads from acinar, alpha and beta cells in scATAC-seq to previously published bulk ATAC-seq ^11^, and observed enriched accessibility signals at both gene bodies and promoter regions of the marker genes *CPA1*, *GCG* and *IGF2* in acinar, alpha, and beta cells, respectively, as expected (**Supplemental Figure 3C,D)**. Further comparison of scATAC-seq cell clusters to the independently identified transcriptomic clusters revealed highly specific correlations between scRNA-seq and scATAC-seq (**Supplemental Figure 3E**), confirming the high quality and reliable cell-type clustering of single cell chromatin accessibility profiles.

### The 3D chromatin architecture of the major pancreatic cell types

To define target genes of regulatory chromatin in a lineage-specific manner, we generated 3D chromatin maps of the three major pancreatic cell types using high-resolution Hi-C chromosome conformation capture on FACS-sorted cell subsets from two biological replicates pooled from 5 healthy donors. Each replicate was sequenced to over 2.2 billion reads, and 1.8 billion reads passing quality control metrics per library were used to construct 3D chromatin maps (**Supplemental Table 2**). Contact matrices were strongly correlated between replicates (stratum-adjusted correlation coefficient >0.9; **Supplemental Figure 4A**). The endocrine cell types (alpha and beta) were more similar to each other than to acinar cells in terms of contact frequency, as expected from their developmental origin and functional properties (**Supplemental Figure 4A**). We observed approximately 5,000 topologically associating domains (TADs) with a median size of 360kb for each cell type (**Supplemental Figure 4B,C**). We further called chromatin contacts at 1kb, 2kb and 4kb resolution independently in each cell type using Mustache^12^ and Fit-Hi-C2^13^ from pooled reads for both replicates (**Supplemental Table 3,** see **Methods**). We observed higher than ∼4-fold enrichment of Hi-C contacts for the loop calls with respect to their local background (**Supplemental Figure 4D**), confirming that the samples allowed for high-fidelity chromatin loop identification.

Next, maintaining the highest resolution, we merged the resultant contact calls from all three cell types to create a combined set of 860,565 consensus chromatin contacts (**Supplemental Figure 4E**). The median distance between loop anchors was 318kb, with 15.6% of called contacts connecting chromatin regions more than 1Mb apart. We annotated these chromatin loops to genes and regulatory chromatin by overlapping loop anchors with gene promoters (-1,500bp to +500bp relative to the transcriptional start site [TSS]) and the open chromatin regions we had identified via scATAC-seq in the corresponding cell types. On average, 58% of the chromatin contacts had at least one anchor annotated to open chromatin regions, while 46% of the chromatin contacts mapped to promoters of protein-coding or long noncoding RNA genes (**Supplemental Figure 4F**). Approximately 20% of the chromatin contacts were annotated to gene-to-open-chromatin-region pairs, in which one anchor overlapped with a gene promoter while the other anchor overlapped open chromatin regions from the corresponding cell type. Compared to random gene-to-open-chromatin-region pairs, those connected by chromatin contacts demonstrated significantly higher co-accessibility of gene promoter and open chromatin region (**Supplemental Figure 4G**, two-sided Wilcoxon rank sum test *P*-value < 2.2e-16). Consistent with previous chromatin conformation capture assays^14, 15^, we also observed a larger number of contacts between genes and distal chromatin regions in the cell types in which these genes were specifically expressed (**Figure 1C**), suggesting a quantitative regulatory effect of chromatin contacts on gene expression.

To further explore the putative regulatory impact of chromatin contacts, we mapped loop anchors to previously characterized chroHMM chromatin states, *i.e*. active promoters, poised enhancers, ‘active enhancers’ (as annotated by chromatin marks, but not functionally tested as such), etc., in endocrine and acinar cells^16^. Compared to distance-matched non-significant Hi-C contacts, significant chromatin contacts were enriched for enhancers (acinar: active enhancer 1, odds ratio (OR) =1.68, active enhancer 2, OR=1.54; alpha: active enhancer 1, OR=1.60, active enhancer 2, OR=1.60; beta: active enhancer 1, OR=1.44, active enhancer 2, OR=1.43) and active TSS (acinar: active TSS OR=2.08; alpha: active TSS OR=1.87; beta active TSS OR=1.68), with ∼68% of chromatin contacts mapping to active enhancers and 8% of contacts mapping to active TSS in the corresponding cell types (**Supplemental Figure 4H,I**). Collectively, these results demonstrate that our robust cell purification approach coupled with high-resolution Hi-C successfully generated comprehensive, high-quality 3D regulatory maps that connect regulatory elements to their target genes in the major human pancreatic cell types.

### Cell type-specific A/B compartmentalization distinguishes pancreatic endocrine and exocrine cells

The genome is divided on the megabase scale into two compartments, termed “A” and “B”, with the former associated with open and the latter with closed chromatin^17, 18^. To investigate if A/B compartmentalization contributes to pancreatic endocrine/exocrine differentiation, we estimated A/B compartments in acinar, alpha and beta cells by eigenvector analysis of the genome contact matrix at 40kb resolution after observed/expected frequency normalization (**Figure 2A**). Prior studies^17, 18^ show that A/B compartments exhibit cell-type specificity, and we likewise detected variation between the pancreatic cell types that substantially exceeded technical variation (**Figure 2B**). Endocrine cell types (alpha and beta) were more similar to each other (jaccard coefficient of A-B assignment = 0.62) than to acinar cells (jaccard coefficient of A-B assignment = 0.51, **Figure 2B**). We observed that 16.9% of the genome was differentially compartmentalized between alpha and beta cells, which was significantly lower than the variant compartmentalization between either endocrine cell type and acinar cells (**Figure 2C**). Further examination of differentially compartmentalized regions between cell types revealed that, compared to invariant compartmentalized regions, the genome was significantly less accessible at regions that were annotated to compartment B in one cell type and compartment A in the other cell types, whereas the genome was more open in differential compartments annotated as A (**Figure 2D**, Wilcoxon Rank Sum Test of all pairwise comparisons, *P*-value < 2.1e-277). Similarly, genes that reside in a compartment A in one cell type and in compartment B in another showed reduced expression in the latter cell type, while genes in located in the A compartment showed increased expression as expected (**Figure 2E**, Wilcoxon Rank Sum Test all pairwise comparison p-value < 3.8e-67).

**Figure 2.**
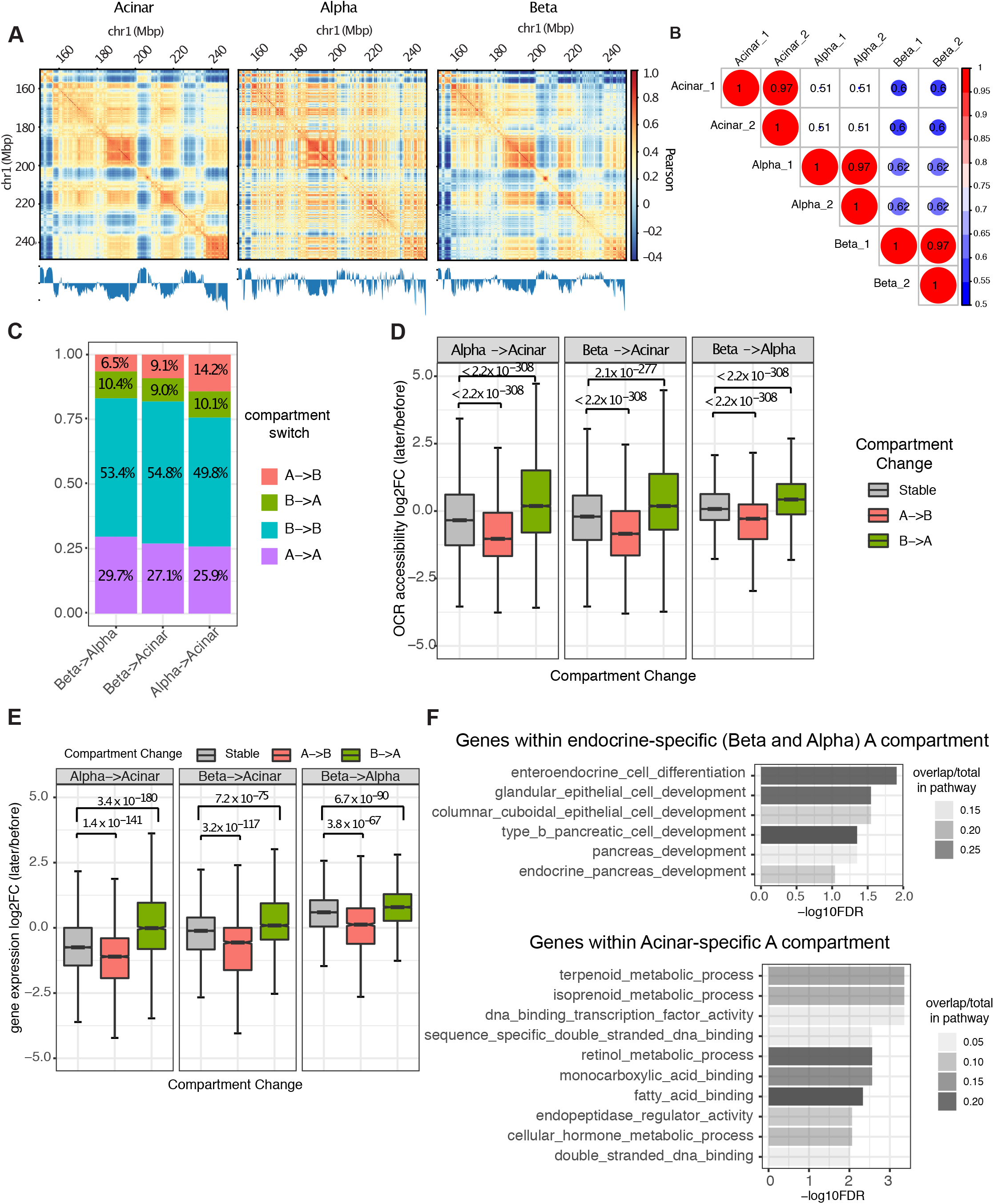
Cell type-specific A/B compartments and their relations to gene expression and regulatory chromatin. **A**, Example of A/B compartment identification using Pearson correlation matrix 40kb resolution on chr1:150,000,000-250,000,000. The plaid pattern suggests chromatin spatially segregates into two compartments. The first eigenvector of the correlation matrix is shown below to derive compartment type with genomic GC content as reference. **B,** Jaccard coefficient of A/B compartment assignment in pairwise comparison. A/B compartment assignment was defined by the sign of first eigenvector for each 40kb bin and compared between Hi-C samples. **C,** Composition of genomic regions that change compartment status or remain the same in pairwise comparison between cell-types. **D and E,** Distribution of fold-change in OCR accessibility (**D**) and gene expression (**E**) at dynamic (A to B or B to A) or stable (A to A or B to B) compartmentalized regions. P value was calculated by two-sided Wilcoxon sum rank test; whiskers correspond to interquartile range. **F**, Pathway enrichment for genes located at endocrine and acinar-specific A compartment. P-values are calculated by hypergeometric test with FDR correction.

When considering acinar cells, we found 763 genes that reside in A compartments that were not present in endocrine cells. These gene sets were significantly enriched for functions associated with the exocrine pancreas such as isoprenoid, terpenoid, and retinol metabolism and endopeptidase regulator activity (**Figure 2F, Supplemental Table 4**). On the other hand, 998 genes that were unambiguously located at A compartments of both alpha and beta cells but not in acinar cells were enriched for endocrine cell differentiation (**Figure 2F, Supplemental Table 4**). Taken together, these results suggest that the differential organization of A and B compartments contribute to shaping the gene expression patterns that drive differentiation of exocrine and endocrine cells in the human pancreas.

### The strength and pattern of chromatin contacts recapitulates cell type-specific gene expression in the human pancreas

To determine how the chromatin contacts that we identified compare between cell types, we performed a quantitative comparison of contact frequency matrices using multiCompareHiC v1.8.0^19^ to identify statistically significant differential looping (FDR < 0.05). Using this approach, we identified 11,052 contacts that were differentially enriched in at least one pairwise comparison (**Supplement Table 5**). These ‘cell type enriched loops’ (CTELs) were further grouped hierarchically into four clusters based on normalized interaction frequencies, with cluster 1 (2,786 loops) representing acinar-specific CTELs, cluster 2 (4,348 loops) representing beta-specific CTELs, cluster 3 (2,378 loops) representing alpha-specific CTELs and cluster 4 (1,540 loops) representing endocrine-shared CTELs (**Figure 3A**).

**Figure 3.**
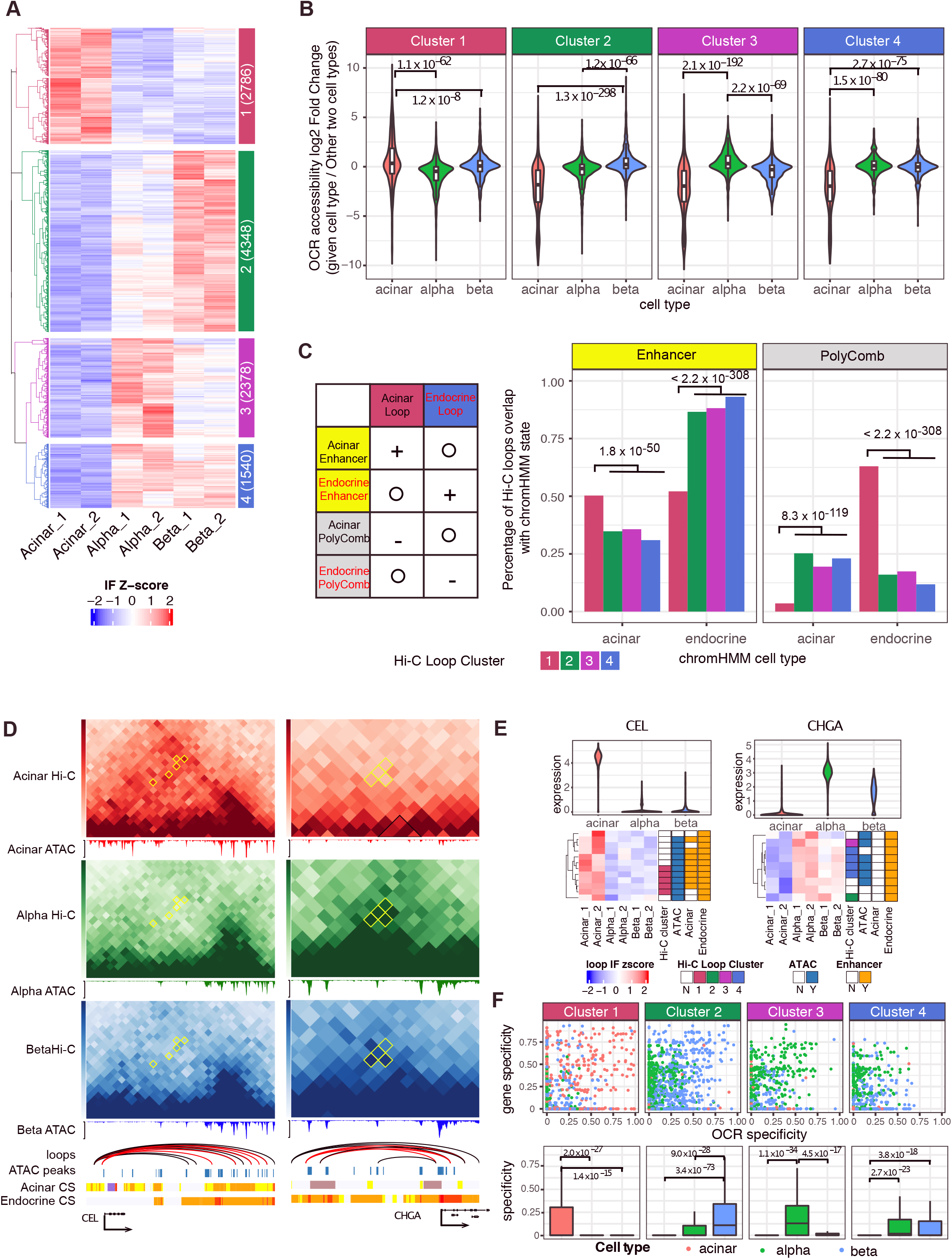
Cell-type differential loops and their effect on gene regulation. **A**, The interaction frequency (IF) heatmap shows distinct clustering of Cell-type enriched loops (CTELs). Each row represents a unique CTEL (see **Methods**) and individual samples are organized in columns; Four clusters are defined by hierarchical clustering. Cluster notion and number of differential chromatin loops are indicated in colored annotation box at left of heatmap. **B**, Distribution of accessibility fold-change of OCRs located at anchors of CTELs. P value is calculated by two-sided Wilcoxon sum rank test; whiskers correspond to interquartile range. **C**, The ratio of clustered CTELs overlapped with enhancer and Polycomb-binding chromatin (PolyComb) from acinar and endocrine cells. Left panel illustrates the expected result based on chromatin enrichment of all chromatin loops in **Supplemental Figure 4I**. Plus sign means “enrichment”, minus sign means “depletion” and circle means “irrelevant”. Right panel shows the observed result in terms of the percentage of clustered CTELs overlapping with given chromatin state. P value is calculated by two-side two sample proportion test. **D**, CTELs connecting promoters of *CEL* and *CHGA* to distal regulatory elements. The Hi-C contact frequency matrix is shown in heatmap with CTDLs in yellow circle. Consensus chromatin loops are represented as an arc with CTELs labelled in red. scATAC-seq is shown as normalized aggregated fragment abundance in each cell type with consensus peaks highlighted in blue bar. Chromatin states (CS) for both acinar and endocrine are represented by a color scheme explained in **Supplemental Figure 4H. E,** The gene regulation scheme on *CEL* and *CHGA*. The interaction frequency (IF) heatmap is shown for all the chromatin loops that connect gene promoter to distal chromatin, with CTELs annotated with corresponding cluster color scheme at left. The chromatin loops are also annotated based on whether distal chromatin anchors are overlapped with consensus ATAC-seq peaks and cell-type enhancer chromatin. **F**, Cell-specificity of gene and OCRs connected by CTELs. The specificity of genes and OCRs in each cell type are calculated using scRNA-seq and scATAC-seq data (see **Methods**). The overall specificity of gene-OCR pair is represented by geometric mean of specificity values of gene and plotted in boxplot. *P*-value is calculated by two-sided Wilcoxon sum rank test; whiskers correspond to interquartile range.

To determine if cell type-specific chromatin contacts are likewise associated with cell type-specific enrichment of regulatory chromatin, we first measured the accessibility of open chromatin regions that overlapped CTEL anchors. This analysis demonstrated that genomic regions with higher interaction frequencies showed increased chromatin accessibility in a given cell type. For example, acinar-specific CTELs connected open chromatin regions that were more open in acinar than in alpha or beta cells (**Figure 3B**). In addition, a comparison with chromHMM-identified chromatin states revealed that acinar-specific CTEL anchors were preferentially enriched for acinar enhancers and, conversely, depleted of acinar polycomb-repressed chromatin, while endocrine-specific CTEL anchors (clusters 2 to 4) were enriched for endocrine enhancers and depleted of endocrine polycomb-repressed chromatin (**Figure 3C**, two-sided two sample proportion test, all *P*-values < 1.8e-50). These results indicate that cell type-enriched chromatin contacts preferentially connect active regulatory regions and recapitulate cell type-specific regulatory signatures.

To explore how cell type-specific chromatin architecture affects gene expression, we focused on signature genes whose promoters were linked to distal regulatory genomic regions by CTELs. For example, *CEL* encodes a lipase secreted from the pancreas into the digestive tract^20^; it was both highly expressed in acinar cells and linked by five acinar-enriched CTELs to distal genomic regions enriched for regulatory chromatin signatures (**Figure 3D, E**). Although these distal regulatory regions were annotated as enhancers in both acinar and endocrine cell types, the enhancer-gene contacts were formed more frequently in acinar than endocrine cells, suggesting that increased physical contact between the *CEL* promoter and distal enhancers drives the gene expression difference between the cell types. Another example is *CHGA,* which encodes chromogranin A, a critical component of secretory vesicles of endocrine cells^21^. Here, six out of nine *CHGA* promoter-connected contacts were found to be specific to endocrine cells, and all distal interacting regions corresponded to endocrine-specific enhancers (**Figure 3D, E**). These findings establish that quantitatively identified CTELs are enriched for cell type-specific enhancers regions.

To globally evaluate the relationship between gene expression, distal *cis*-regulatory element activity, and physical gene-to-*cis*-regulatory-element contacts, we focused on the 2,910 (26.3%) CTELs that we could annotate to gene-open chromatin region pairs, and derived the gene expression and open chromatin region accessibility specificity weight matrix from our single cell scRNA-seq and scATAC-seq data using CELLEX^22^ (see **Methods**). We found that gene-open chromatin region pairs with high cell type-specificity were preferentially joined to contacts enriched in the same cell type (**Figure 3F**), i.e. acinar-specific CTELs (cluster 1) connect open chromatin regions and genes with high specificity in acinar cells, while beta-specific CTELs (cluster 2) connect open chromatin regions and genes with high specificity values in beta cells (**Figure 3F**). Overall, these findings suggest that differential chromatin contacts play an important role in cell type-specific regulation of gene expression in the pancreas.

### Cell type-specific transcription factor accessibility and promoter connectivity cooperate to establish lineage-specific gene expression programs

Changes in chromatin structure at genomic regions are generally accompanied by dynamic binding of transcription factors^23, 24^. The development and differentiation of the pancreas is governed by the coordinated action of lineage-specific transcription factors that direct cell-selective transcriptional programs^25–30^. To investigate which transcription factors are enriched in cell type-specific regulatory chromatin organization in the human pancreas, we employed chromVAR^31^ to assess the global accessibility of transcription factor motifs from the JASPAR 2020 database^32^ in the data derived from scATAC-seq. Motif enrichment in each cell type was represented by an accessibility deviation score reflecting the accessibility of peak sets in the selected cell type relative to the basal assumption of equal chromatin accessibility across all cells. **Figure 4A** summarizes these data, which confirm previously known motif enrichment patterns. For example, the motif occupied by the endocrine transcription factor NEUROD1 was enriched in alpha, beta, delta and epsilon cells, while the motif preferred by NKX6-1, PDX1 and PAX4 was enriched in beta and delta cells. Additionally, the binding sites of the exocrine transcription factors NR5A2, GATA4 and GATA6 were enriched in ductal and acinar cells (**Figure 4A, B**). Further clustering of cell types based on motif enrichment revealed close relationships between acinar and ductal (Pearson correlation = 0.78), endothelial and mesenchyme (Pearson correlation = 0.77), alpha and epsilon (Pearson correlation = 0.8), and beta and delta cells (Pearson correlation = 0.64, **Figure 4C**). These findings are consistent with the transcriptome profiles of these cell types measured by scRNA-seq (**Supplemental Figure 2D**), suggesting that cell lineage-enriched transcription factors together with differential regulatory chromatin accessibility change across cell types to determine cell-specific gene expression.

**Figure 4.**
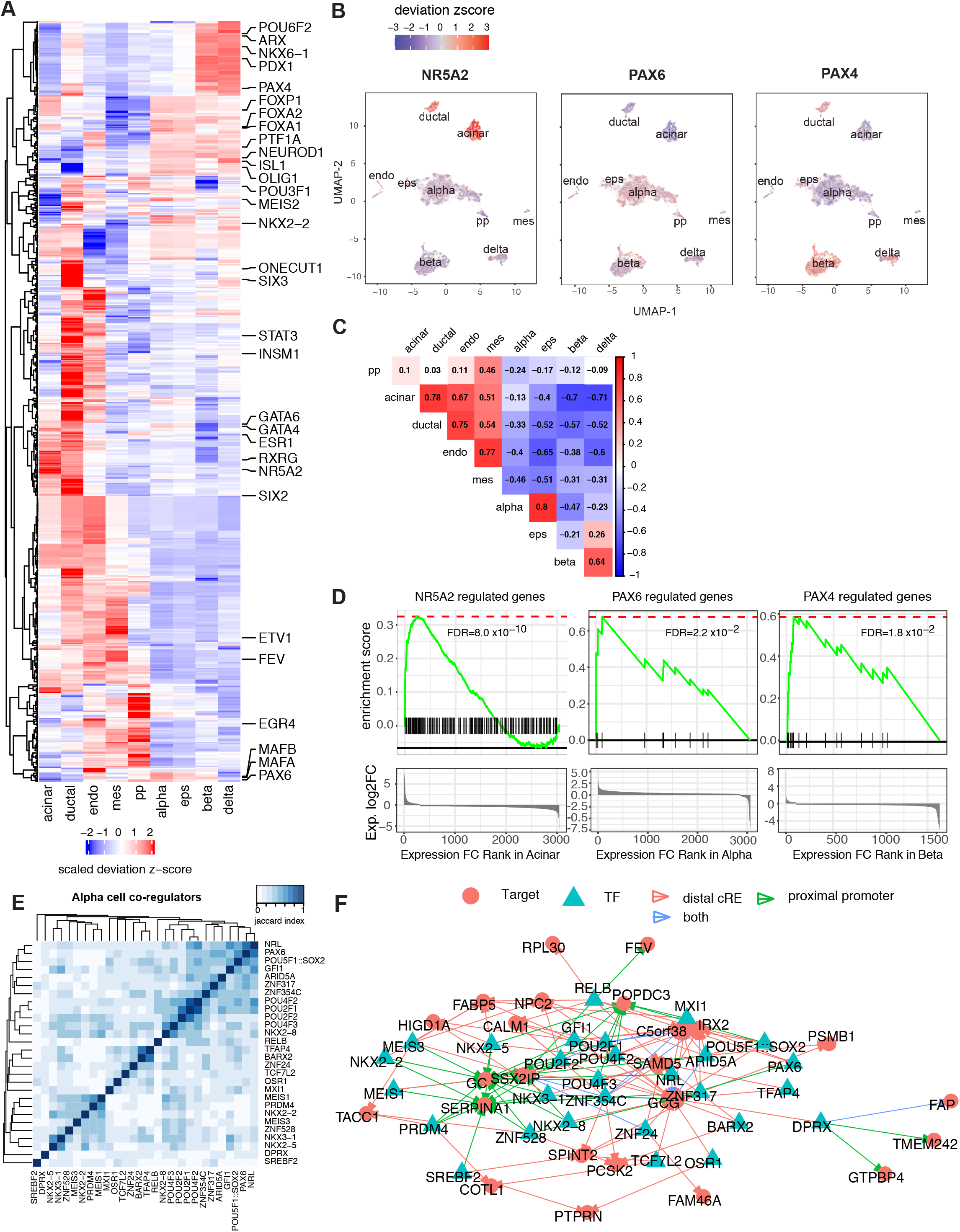
Cell-type specific transcription factor (TF) motif enrichment and TF-gene network. **A**, chromVAR motif enrichment for 438 cell-type-specific TFs. The identification of cell-type-specific TFs was explained in **Methods** and full list is in **Supplemental Table 6**. The enrichment was measured by deviation z-score. **B**, The enrichment for NR5A2, PAX6 and PAX4 motifs for each cell projected onto UMAP. **C**, Pearson correlation of TF enrichment between cell types. The deviation z-score was used for correlation. **D**, GSEA pre-rank enrichment of cell-specific expressed genes by the NR5A2, PAX6 and PAX4 downstream genes. Genes are ranked by gene expression log2FC in the given cell type from highest to the lowest. Gene sets are defined by TF downstream genes of which either promoters or distal cis-regulatory elements contain the given TF motif. Details of enrichment is in **Supplemental Table 7**. **E**, Association index analysis reveals alpha-specific TF modules. TF sets are defined cell-specific TFs with significant GSEA enrichment (**Supplemental Table 7**). Matrices show the Jaccard similarity between TF pairs based on occurrence of co-regulated downstream gene. **F**, TF-gene network for alpha-specific TF sets. The nodes are either TF (green triangle) or downstream genes (red circle) from leading edge in **Supplemental Table 7**. The edges represent how TF and downstream gene connected: TF motif at only distal cRE (red), TF motif at only gene promoter (green) and TF motif at both gene promoter and connected cREs (blue).

To further explore this concept, we mapped regions harboring accessible transcription factor motifs to the genes they likely regulate via physical contact in acinar, alpha and beta cells using our genome-scale chromatin connectivity datasets. Focusing on transcription factor binding sites in cell type-specific open chromatin (see **Methods**), we identified between 23 and 2,179 (mean 580) likely downstream genes per transcription factor motif. Among all 438 TF motifs exhibiting cell type-specific accessibility (**Supplemental Table 6,** see **Methods**), 65% (283) were physically connected to genes with significantly higher expression in that same cell type compared to other cell types (**Supplemental Table 7**). For example, NR5A2, whose motif was specifically enriched in the open chromatin landscape of acinar and ductal cells (**Figure 4A, B**), was predicted to regulate 138 downstream genes through its binding to promoters or promoter-connected distal regions. These 138 genes, including *CPA1*, *CTRB2*, *CTRB1* and *PLA2G1B*, were enriched for high expression in acinar cells compared to other cell types (GSEA pre-rank permutation test FDR = 8.0e-10, **Figure 4D, Supplemental Table 7**). Similarly, PAX6 and PAX4, whose motifs were specifically enriched in endocrine cells, were localized to downstream genes with elevated expression in alpha and beta cells, respectively (GSEA pre-rank permutation test PAX6: FDR = 0.02; PAX4: FDR=0.018, **Figure 4D, Supplemental Table 7**).

Next, we calculated the binding similarities between transcription factors to define jointly acting regulators in a cell-specific manner, based on their shared downstream target genes (**Supplemental Table 7**). As expected, TFs with similar motif positional weight matrices clustered together (e.g. POU class 2 and class 4 TF motifs in alpha co-regulators; **Figure 4E**, KLF family TF motifs among acinar co-regulators **Supplemental Figure 5A**). However, we also found TFs with divergent motifs that grouped together because they target similar sets of downstream genes. For example, NRL, PAX6 and POU5F1/SOX2 motifs are highly colocalized with alpha cell open chromatin, all having motifs in the alpha-specific genes *GCG*, *IRX2* and *C5orf38* (**Figure 4E, Supplemental Table 7**). Similarly, NKX6−1 and POU6F1 motifs, which are enriched in beta cell open chromatin, are each connected to the *MAFA, HADH, ADCYAP1, SYNE2, SORL1, GNAS, SAMD11* and *SCD5* genes that are likewise highly expressed in beta cells (**Supplemental Figure 5B, Supplemental Table 7**). In addition, we were able to identify cell-specific “hub” regulators based on the number of targeted downstream genes and their involvement in the cohesiveness of TF-target gene transcription regulatory network. In alpha cells, POU2F1 and RELB motifs targeted 8 alpha-specific genes that showed elevated expression in alpha cells (**Figure 4F**, **Supplemental Table 6 and 7**). In beta cells, NHLH2 and ARID3A motifs targeted 14 and 25 downstream genes, respectively, that were specifically expressed in beta cells, and *NHLH2* itself showed elevated expression in beta cells (**Supplemental Figure 5C**, **Supplemental Table 6 and 7**). This integrated analysis of the transcriptome, open chromatin landscape, and nuclear architecture of the major cell types of the human pancreas refines our understanding of lineage-specific regulators that operate to establish pancreatic gene regulatory networks.

### Cell type-specific assignment of diabetes-associated variants to their target effector genes

Genetic variants associated with diabetes and diabetes-relevant traits are enriched at chromatin accessible sites in various pancreatic islet cell types^33^. However, our current 3D epigenomic analyses of purified human pancreatic alpha, beta, and acinar cells allows a previously unattainable, direct view of specific genes that these T2D variants target in the major cell types of the pancreas. To achieve this goal, we focused on accessible chromatin regions that were connected to at least one target gene through either long-range chromatin conformation or promoter proximity, and performed partitioned heritability linkage disequilibrium (LD) score regression (LDSC) ^34^ on type 1 and 2 diabetes^33, 35^, glycemic traits^36^ and other complex traits^37–40^ independently in acinar, alpha and beta cells. Overall, we found significant enrichment of fasting glucose (alpha: enrichment=23.8, p-value=1.1e-6; beta: enrichment=35.0, *P*-value=2.3e-5), T2D (alpha: enrichment=13.8, p-value=3.3e-3; beta: enrichment=17.1, *P*-value=2.4e-4), sleep duration (beta: enrichment=7.6, p-value=6.9e-5) and chronotype (alpha: enrichment=8.6, *P*-value=3.9e-6; beta: enrichment=10.0, *P*-value=2.7e-7) in beta and alpha cells, and enrichment of pancreatic cancer (acinar: enrichment=5379, *P*-value=1.6e-3) in acinar cells (FDR < 0.1, **Figure 5A, Supplemental Table 8)**. We also observed additional significant enrichment of fasting glucose (enrichment=17.3, *P*-value=3.0e-3) and a more nominal enrichment of T2D (enrichment=14.4, *P*-value=7.5e-3) in acinar cells, which implicates epigenetic chromatin-structure-based connections between pancreatic exocrine cells and T2D.

**Figure 5.**
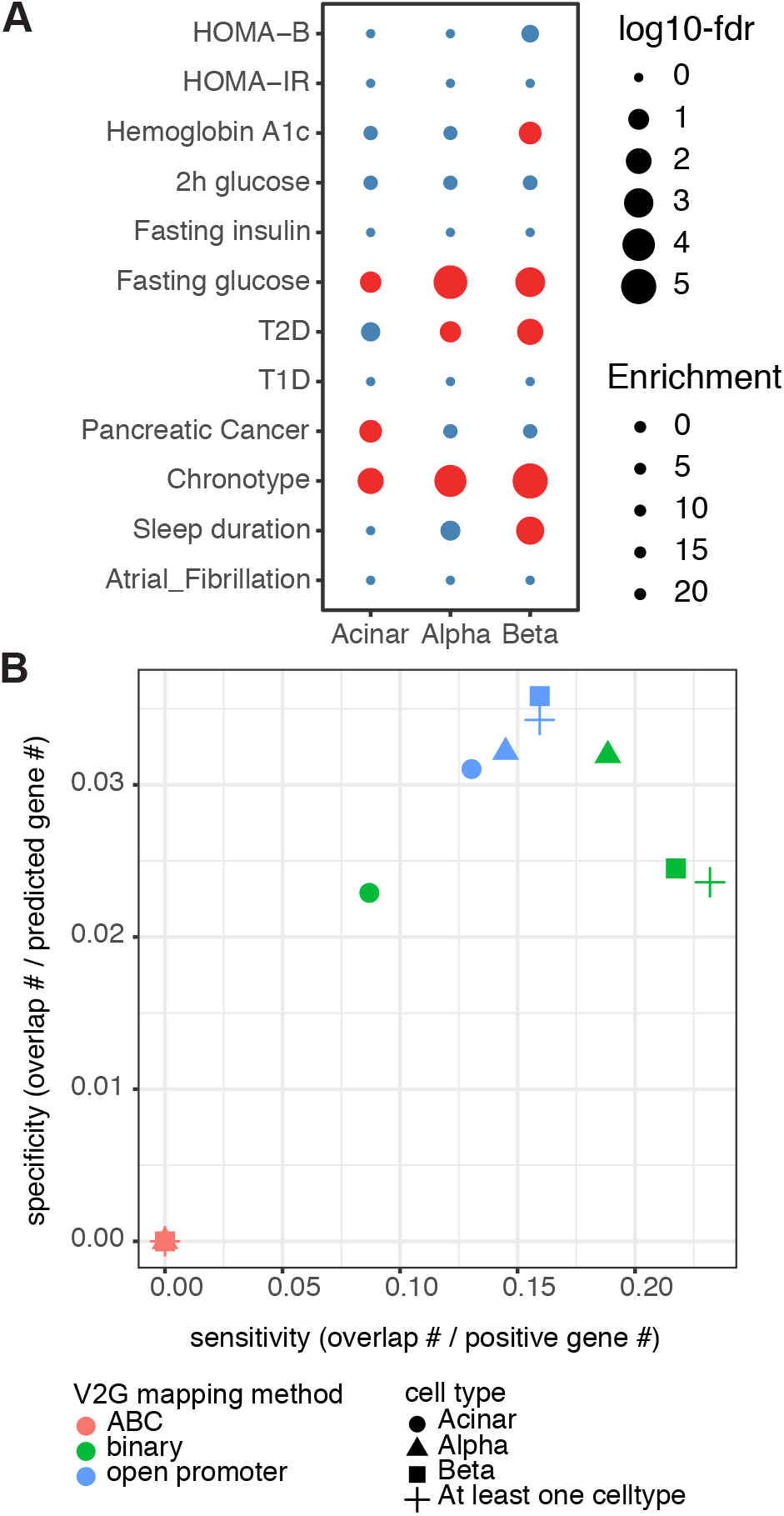
Chromatin contacts link T2D risk variants to target genes with cell-type stratification. **A**, Partitioned LD score regression enrichment of gene-connected regulatory chromatin for T2D and relevant traits. Significant enrichment FDR < 0.1 is indicated with red. **B**, Pathway enrichment of putative effector genes predicted by chromatin-guided variant-to-gene mapping. P-values are calculated by hypergeometric test with FDR correction. **C**, Posterior probability of putative causal variants. 352 putative causal variants were defined either by chromatin-guided variant-to-gene mapping or randomly picked from distance-matched credible sets for T2D sentinel signals. *P*-values are calculated by two-sided Wilcoxon rank sum test.

A number of approaches have been developed to assign variants to genes by leveraging regulatory chromatin landscape and/or nuclear architectural data. Here we compared two general approaches: (1) The Activity-by-Contact (ABC) model^41^, which computes a score that predicts gene-enhancer connections by combining the read counts of enhancer accessibility (ATAC-seq or DNAse-seq) and presumed activity (H3K27ac ChIP-seq) with the normalized contact frequency (HiC) between putative enhancers and gene promoters, or (2) binary calls using the intersection of significant cell-specific loops called from our Hi-C data and *cis*-regulatory elements defined by ATAC-seq with corresponding gene promoters (**Supplemental Figure 6,** see **Methods**). Notably, it was previously reported that an averaged HiC matrix of several cell lines performed similarly to cell type matched HiC data in the ABC model^41, 42^. Using our cell-type-specific Hi-C contact frequencies in acinar, alpha and beta, the ABC model linked 12,114 putative enhancers to 9,062 target genes (31,103 enhancer-gene pairs) in all three cell types (ABC score >= 0.02, **Supplemental Figure 7A,** <u>Gene Cis-element Interactome in Pancreas Hi-C shinyApp</u>). In contrast, our chromatin loop binary calls identified four times as many gene-distal element connections, with 146,472 *cis*-regulatory-element-gene pairs associating 55,003 open chromatin regions to 20,029 genes in at least one of the three cell types (**Supplemental Figure 7A,** <u>Gene Cis-element Interactome in Pancreas Hi-C shinyApp</u>).

Even though our chromatin loop binary calls did not constrain the *cis*-regulatory-elements to open chromatin regions marked by the H3K27ac signal like the ABC model does, we observed that on average fewer than 10% of the gene to putative enhancer pairs identified by the ABC model were also present in the binary calls (**Supplemental Figure 7A**). In order to evaluate how the two different approaches affect variant-to-gene mapping, we applied the two gene to distal-element maps to T2D GWAS signals^35^. The currently known 403 sentinel SNPs for T2D (**Supplemental Table 9**) correspond to 8,859 proxy SNPs that are within close linkage disequlibrium (LD, r^2^ > 0.8) using the 1,000 genome European population as the reference and 38,758 fine-mapped variants of which 5,768 variants are shared between proxies (R2 > 0.8) and 95% credible set. The chromatin loops we mapped above in the major pancreatic cell types revealed contacts related to 165 sentinels, for which we could map 1162 open SNPs to 922 distally located gene promoters. Limiting to just the 165 sentinel SNPs for which our pancreas-specific Hi-C maps and ATAC-seq data were informative, our experimental approach narrowed down the list of possible causal SNPs from 34,993 to 1162, *i.e.* a >96% reduction of the search space for causal variants.

In contrast, the ABC model linked only 301 putative causal variants (corresponding to 96 sentinel SNPs) to 411 gene promoters in our three cell types (**Supplemental Table 10, Supplemental Figure 7B**). Although 56.8% of the proxy SNPs identified by the ABC model were also implicated by our chromatin loop binary approach, the two methods linked SNPs to very different sets of genes, with only 42% of the genes (8.4% proxy-gene pairs) from the ABC model recapitulated by the chromatin loop binary approach (**Supplemental Figure 7B**).

Next, we compared the implicated genes to a set of independently curated T2D positive control genes, which include (1) genes whose perturbation causes a Mendelian form of diabetes (2) genes whose encoded protein acts as a drug target for the common disease, or (3) coding variants discovered in case-control exome studies of T2D^43^. None of the genes identified by the ABC model overlapped with these T2D positive control genes, while our chromatin loop binary calls detected 23.2% of positive control gene set with various sensitivity and specificity among the different cell types (**Figure 5B**). Given these observations, we elected to use the gene to distal-element map identified by our chromatin loop binary approach for variant-to-gene mapping and complimented the list of putative causal variants with open proxies located at gene promoters we had identified (**Figure 5B, Supplemental Table 10**).

Finally, we found that 194 of the 403 T2D GWAS sentinels (32%) were in LD with at least one variant situated in an open gene promoter (147 sentinels) or a distal OCR found in direct contact with a gene promoter (165 sentinels) in the three human pancreatic cell types analyzed. This represents 1,388 putative casual variants mapping to a total of 1,091 genes across acinar, alpha and beta cells at these loci (**Supplemental Table 10**). Out of 4,750 casual-variant-gene pairs identified by our method, only 369 (7.8%) were reproduced by previously islet promoter-enhancer promoter capture Hi-C^9^. Conversely, of the prior whole islet variant-gene pairs reproduced here, 98.4% (363) are physically connected by Hi-C loops in alpha and beta cells, as expected. Further evaluation of a variant’s impact on the immediate surrounding genomic location found that 869 proxies (57.5%) show a strong disruptive effect on at least one TF binding site (**Supplemental Table 11**).

Importantly, in addition to dramatically reducing the number of likely putative causal variants for T2D, our analyses mapped each of the 194 T2D risk loci containing at least one putative causal-variant-gene pair to their likely effector transcripts. In total, we provide variant-to gene-to cell type mapping for 900 genes, many of which previously were either not supported, or only scantly supported, as T2D relevant (**Supplemental Table 12**), which will greatly facilitate functional interrogation of these gene in the future. To further dissect the cell-specificity of T2D variant-to-gene connections, we examined cell type-specific gene expression, variant-containing open chromatin region accessibility, and gene-open chromatin region contact frequency for each putative causal-variant-gene pair. We discovered 16 putative effector genes that, together with their interacting variant-containing open chromatin regions, demonstrated high cell-type specificity (specificity weight >=0.5, **Figure A,B**). As expected, most of cell-type specific putative effector genes, e.g., *INS*, *DGKB*, and *P2RY1*, were specific to insulin-producing beta cells (**Figure 6B**). However, we also implicated 5 putative effector genes likely to function in non-beta cells; the alpha cell-enriched *WFS1* encodes Wolframin, a transmembrane protein regulating cellular Ca^2+^ homeostasis, and the acinar-specific gene *KCNQ1*, *SLC39A8*, *CTRB1*, and *CPA4* (**Figure 6B**). Importantly, with the exception of *TRPM5*, all chromatin interactions of these causal-variant-gene pairs demonstrated higher contact frequency in their specific cell type of action (**Figure 6B**), suggesting the importance of chromatin looping in the cell-type specific regulation of T2D pathogenesis. For example, at the *TH* locus (rs4929965), a beta-cell specific open chromatin region harboring three putative causal variants (rs7482891 A>G r^2^=0.98, rs4929964 T>G r^2^=0.98 and rs4929965 A>G r^2^=1), physically contacted the *INS* promoter over 12kb away preferentially in beta cells (**Figure 6C)**. The same open chromatin region also interacted with the *TRPM5* promoter over 250kbp downstream in both alpha and beta cells, with a higher contact frequency in alpha cells (**Figure 6C, Supplemental Table 10**). Further examination of the three putative causal variants revealed that all of them resided at transcriptional factor binding sites (rs4929965 A>G: KLF4, ZNF148, MAZ and PLAGL2; rs4929964 T>G: NR2C2; rs7482891 A>G: KLF12 and GLIS1) and were predicted to strongly disrupt factor occupancy (**Supplemental Figure 8A, Supplemental Table 11)**. Similarly, at the *WFS1* locus (rs10937721), a putative endocrine-specific causal proxy variant (rs4234731 A>G r^2^=0.88) in intron 7 of *WFS1* physically contacted the *WFS1* promoter in both alpha and beta cells, with an additional alpha-cell specific contact further upstream (**Figure 6D**). Importantly, this variant is predicted to strongly disrupt the binding of the endocrine transcription factor NEUROD2 (**Supplemental Figure 8B**). Numerous publications have associated with *WFS1 locus* with Mendelian and common forms of diabetes^34, 35^. A mouse study found that disruption of *WFS1* gene causes progressive beta cell loss and impairs insulin exocytosis^36^. However, our chromatin looping approach found the connection between the putative causal variant rs4234731 and *WFS1* promoter is more specific to alpha cells; elevated gene expression, increased variant accessibility, and higher contact frequency are all observed in in alpha cells compared to beta cells (**Figure 6B**), necessitating the future investigation of *WFS1* function in the alpha cell context.

**Figure 6.**
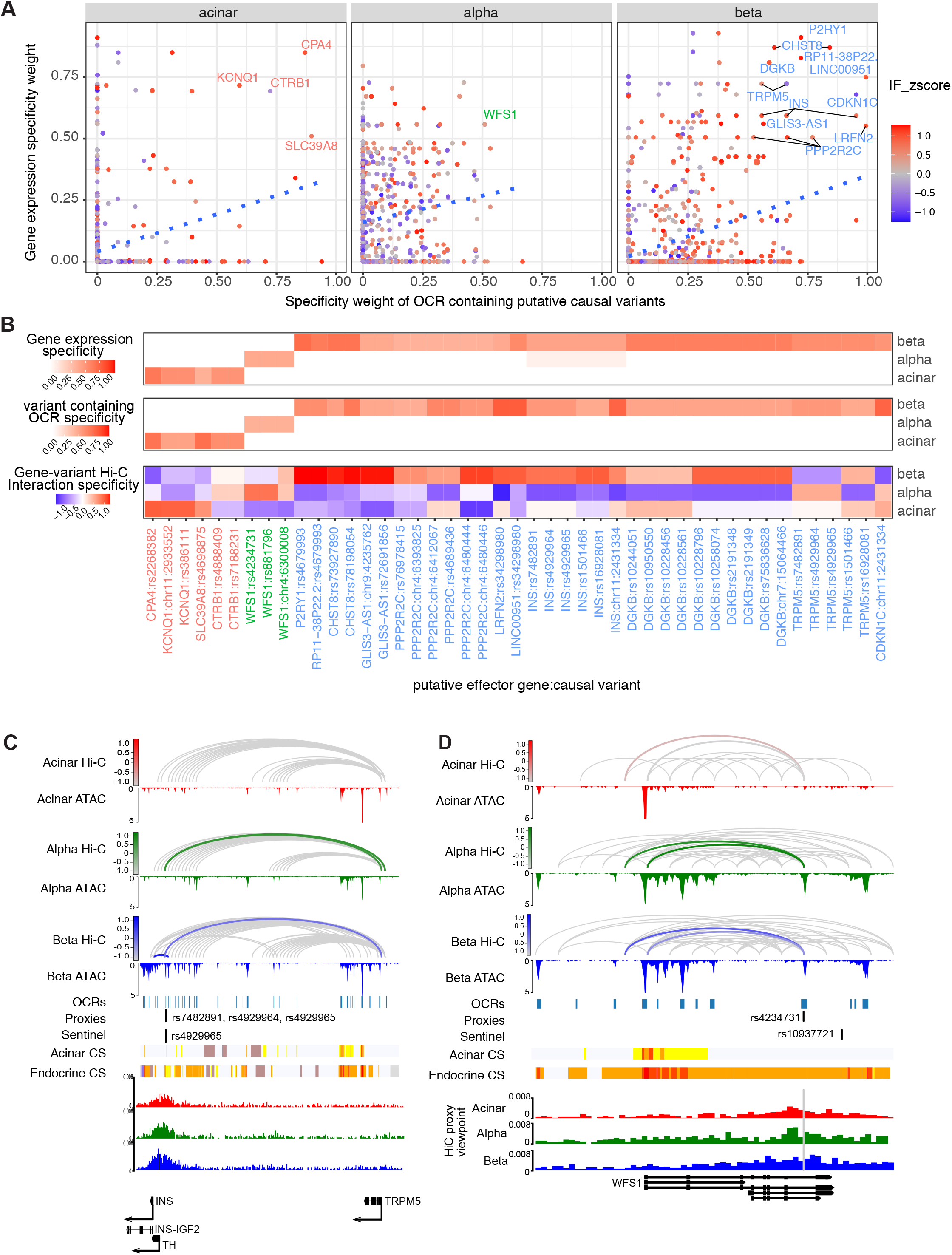
Chromatin contacts link T2D risk variants to target genes with cell-type stratification. **A**, Cell-specificity of putative causal variants and their target gene. The variant-containing gene-OCR pairs with high cell-specificity (specificity >= 0.5) were labelled with target gene name for each cell type **B**, The gene-causal-variant pairs were clustered based on cell-type specificity **C**, Cell type-specific chromatin looping between causal variants (rs7482891, rs4929964 and rs4929965) and target genes *INS* and *TRPM5* and the T2D *TH* (rs4929965) locus. The chromatin loops that connect causal variants and target genes are highlighted with cell-type specific color (acinar: red, alpha: green and beta: blue), with hue scale indicating Hi-C interaction frequency. **D**. TF motifs disrupted by causal variants (rs7482891, rs4929964 and rs4929965) at the T2D *TH* (rs4929965) locus. **E**, Cell-specific chromatin looping between variant rs4234731 and the target gene *WFS1* at the T2D rs10937721 locus. The chromatin loops that connect putative causal variants and target genes are highlighted with cell-type specific color (acinar: red, alpha: green and beta: blue) with the hue scale indicating the Hi-C interaction frequency. **F**, TF motifs disrupted by t variants rs4234731 at the T2D *WFS1* locus.

To further prioritize causal genes and elucidate mechanisms involved in T2D, we compared the variant-gene pairs predicted for T2D to our analyses of other relevant traits (**Supplemental Table 13**), including fasting glucose (FG), Hemoglobin A1C (HbA1c), fasting insulin (FI), 2h glucose (2hGluc) and type 1 diabetes (T1D). Among 4,750 putative causal gene-variant pairs in T2D, 241 pairs (5.1%) were identified within one or more other relevant trait (**Supplemental Figure 9A**) and correspond to 23 T2D sentinel signals. Although T1D and T2D shared most gene-variant pairs, 15 out of 16 signals that show statistically significant multi-trait genetic etiology with T2D were only colocalized with glycemic traits (see **Methods, Supplemental Table 14**), suggesting pleiotropic effects of genetic variants mainly between T2D and glycemic traits. Next, we employed colocalization with glycemic traits to infer potential mechanisms for causal variants in T2D. For example, fasting glucose and hemoglobin A1C were colocalized with T2D at sentinel signal rs10228066 with a posterior probability of 0.819 (**Supplemental Figure 9B**). Variant rs10228796, which connects specifically in beta cells to the promoter of diacylglycerol kinase (*DGKB*), (**Figure 6B**), explained 33.6% of this colocalization by variant to gene-mapping (**Supplemental Figure 9C**), implicating the beta-cell specific chromatin connection rs10228796-*DGKB* in genetically contributing to the elevated levels of fasting glucose and hemoglobin A1C observed in T2D patients. Mechanistically, the inhibition of diacylglycerol kinase reduces beta cell insulin secretion and leads to disruption of peripheral glucose metabolism^44^. In addition, defective glucose homeostasis increases synthesis of diacylglycerols which leads to lipo-toxicity of the beta cells, resulting beta cell dysfunction in T2D^45^. This example of prioritizing the rs10228796-*DGKB* interaction by both multi-trait variant-to-gene-mapping and genetic colocalization data demonstrates that our pancreatic cell chromatin architecture-guided variant-to-gene mapping not only sheds light on the molecular mechanistic effect of causal variants on associations among genetic-relevant traits, but also infers the specific cell-type context for further functional study on the relationship between these traits.

## Discussion

T2D is a major health problem and an immense economic burden on our health care system. Due to the sheer prevalence of type 2 diabetes, this disease has received widespread attention by human geneticists and has often led the way in how to best tackle the discovery of loci, and has thus been the focus of more GWAS analyses than any other disorder studied to date. The GWAS approach has identified hundreds of loci that are associated with T2D disease risk with high statistical significance that have been reproduced over multiple patient cohorts. Unfortunately, given the complexity of gene regulation in humans, in many cases the effector transcript and cell type of action of a given locus are uncertain. Of note, the GWAS approach was never designed with the explicit intention of interrogating the actual underlying causal variant conferring increased risk for disease, but only to ‘tag’ the approximate location of a disease variant typically down to a few hundreds of kilobases^46–48^. Thus, for almost all signals, the effector transcript and the cell type(s) of action of the relevant gene are unknown. However, the identification of the effector transcript and site of action is needed in order for us to understand better the etiology of T2D and to identify novel drug targets.

In this study, we addressed the key issues of causality at key T2D GWAS-implicated loci in order to identify ‘culprit’ genes. We tackled this challenge through the integration of GWAS findings with scATAC-seq, scRNA-seq and high-resolution Hi-C analyses from sorted key cell types of the human pancreas, allowing us to map variant-to-gene-by-cell type relationships for 900 genes (Supplemental Table 12), thus delivering critical information required for testing and evaluating gene function in individual pancreatic cell types. This complements the previously variant-to-gene mapping work by combining these state-of-the-art techniques to drill down cell-type specific causal effector genes within pancreas for the key loci established in T2D GWAS efforts. This in turn offers the eventual potential to uncover corresponding diagnostically actionable variants harbored within these genes to better serve precision medicine in the T2D field going forward.

With the advantage of chromatin conformation maps established in finely sorted pancreatic cell types, our study also revealed that not all T2D loci are likely to act within the insulin-producing beta cell. For example, we discovered that the Wolfram syndrome 1 (*WSF1*) gene, previously shown to be critical for beta cell survival in mice^49, 50^, is likely to function additionally in glucagon-producing alpha cells in humans (**Figure 6**). While further experimental validation is required, this finding suggests that caution is warranted when inferring phenotypes of null alleles in mice to those of weak enhancer mutations that most T2D risk variants represent. Perhaps even more surprising is our discovery of that the T2D risk locus tagged by rs2268382 connects to *CPA4*, encoding Carboxypeptidase 4 (CPA4) specifically in acinar cells, suggesting the exocrine pancreas as the likely tissue of action for this T2D risk locus. In support of this finding, exocrine pancreas insufficiency is frequently associated with diabetes and has been suggested as a critical factor in the development of the disease ^51^. In sum, our integration of GWAS findings with cell type-specific high-resolution Hi-C, single cell ATAC-seq and transcriptome data has allowed for the identification of hundreds of likely variant-to-gene pairs and will serve as a crucial resource for the diabetes research community.

## Materials and Methods

### Experimental model and subject details

Pancreatic islets were procured by the HPAP consortium under the Human Islet Research Network (https://hirnetwork.org/) with approval from the University of Florida Institutional Review Board (IRB # 201600029) and the United Network for Organ Sharing (UNOS). A legal representative for each donor provided informed consent prior to organ retrieval. Organs were recovered and processed as previously described^52^. Supplemental Table 1 summarizes donor information. Pancreatic islets were cultured and dissociated into single cells as described^11^.

Total dissociated cells were subjected to FACS sorting as reported previously^53^.

### Hi-C library preparation

Hi-C library preparation on FACS-sorted pancreatic cells was performed using the Arima-HiC kit (Arima Genomics Inc; cat# A410030), according to the manufacturer’s protocols. Briefly, cells were crosslinked using formaldehyde. Crosslinked cells were then subject to the Arima-HiC protocol, which utilizes multiple restriction enzymes to digest chromatin. Arima-HiC sequencing libraries were prepared by first shearing purified proximally-ligated DNA and then size-selecting 200-600 bp DNA fragments using AmpureXP beads (Beckman Coulter; cat# A63882). The size-selected fragments were then enriched using Enrichment Beads (provided in the Arima-HiC kit), and then converted into Illumina-compatible sequencing libraries with the Swift Accel-NGS 2S Plus DNA Library Kit (Swift, 21024) and Swift 2S Indexing Kit (Swift, 26148). The purified, PCR-amplified DNA underwent standard QC (qPCR, Bioanalyzer, and KAPA Library Quantification [Roche, KK4824]) and was sequenced with unique single indexes on the Illumina NovaSeq 6000 Sequencing System at 2x101 base pairs.

### scRNA-seq data generation and analysis

The Single Cell 3’ Reagent Kit v2 or v3 was used for generating scRNA-seq data. 3,000 cells were targeted for recovery per donor. All libraries were validated for quality and size distribution using a BioAnalyzer 2100 (Agilent) and quantified using Kapa (Illumina). For samples prepared using ‘The Single Cell 3’ Reagent Kit v2’, the following chemistry was performed on an Illumina HiSeq4000: Read 1: 26 cycles, i7 Index: 8 cycles, i5 index: 0 cycles, and Read 2: 98 cycles. For samples prepared using ‘The Single Cell 3’ Reagent Kit v3’, the following chemistry was performed on an Illumina HiSeq 4000: Read 1: 28 cycles, i7 Index: 8 cycles, i5 index: 0 cycles, and Read 2: 91 cycles. Raw sequencing reads were aligned to the hg19 reference genome and quantified using CellRanger v3.1.0 (http://10xgenomics.com). Seurat R package v3.0 ^54^ was used for data quality control, preprocessing and dimensional reduction analysis. After gene-cell data matrix generation for different donors, matrices were merged and poor-quality cells with < 200 expressed genes and mitochondrial gene percentages > 10% were excluded.

### scATAC-seq data generation and analysis

Handpicked islets (approximately 5,000 IEQs) were transferred into a 15 mL conical tube and 10 mL of 1xPBS w/o Ca^2+^, Mg^2+^ (Rockland, MB-008) added. Islets were centrifuged for 2 min at RT, 180 x g and the supernatant removed. Islets were then incubated with 1 mL of warm (37 °C) 0.05% Trypsin (Invitrogen, 25300054) at 37 °C for 9 min with tituration with a 1 ml pipette at 0, 2, 4 and 7 min. The digestion was stopped by addition of 1 mL of 100% FBS (Hyclone, SH3091003) and dissociated islets passed through a BD FACs tube with strainer top (Corning 352235). After rinsing with 1 mL of 100% FBS and transfer to a 15 mL conical cells were centrifuged for 4 min at 400 x g. Cell pellets were resuspended in 1 ml PBS with 10% FBS and centrifuged for 4 min, 400 x g two times before the cells were counted after resuspension in 0.04% BSA in PBS. 5,000 nuclei were processed for scATACseq using Chromium Single Cell ATAC Reagent Kits and sequenced on an Illumina Novaseq 6000 instrument.

Raw fastq files were aligned to the hg19 reference genome and quantified using CellRanger atac v1.2.0 (http://10xgenomics.com). The resulting sorted BAM files were hierarchically structured into hdf5 snapfiles using snaptools v 1.0.0. Quality control was conducted using SnapATAC v.1.0.0^55^ by only keeping the cells with less than 3∼6 logUMI, 10% ∼ 60%promoter ratio. The cells with more than 10% mitochondrial ratio and less than 1000 mapped fragments were also removed. Then, we binarized fragments into 1-kb bins and removed bins on mitochondria, chromosome X, chromosome Y and ambiguous chromosomes, overlapping with the ENCODE blacklist as well as top 5% bins that overlap with invariant features such as the housekeeping gene promoters. Cells with a bin coverage less than 1,000 were excluded.

‘Diffusion map’ was applied as a dimension reduction method using function runDiffusionMaps on 50 eigenvectors and the first 14 eigenvectors were then selected for further batch effect correction using Harmony v.1.0.0^56^ and a k-nearest neighbor graph generation using function runKNN with k = 15. Clustering was performed using the igraph algorithm and UMAP embeddings were generated to visualize the data. We manually assigned each identified cluster to pancreatic cell types based on promoter accessibility of known marker genes.

Fragments originating from cells belonging to the same clusters were pooled using snaptools, and pseudo bulk open chromatin regions were called for each cell type using MACS2 v2.1.2^57^ with the parameters --nomodel --shift 100 --ext 200 --qval 5e-2 -B --SPMR. Differential accessible regions (DARs) across different cell types were next identified using the findDAR function in the SnapATAC package with exactTest and the parameter cluster.neg.method as"knn" and bcv as 0.4. DARs were determined by an FDR < 0.05 and absolute log fold change > 1. Cell-type specific open chromatin regions were DARs that are only significant in that given cell types but not others.

### Hi-C data analysis

Paired-end reads from two replicates were pre-processed using the HICUP pipeline v0.7.4^58^, with bowtie as aligner and hg19 as the reference genome. The alignment .bam file were parsed to .pairs format using pairtools v 0.3.0 and pairix v0.3.7, and eventually converted to pre-binned Hi-C matrix in .cool format by cooler v 0.8.10 with multiple resolutions (500bp, 1kbp, 2kbp, 2.5kbp, 4kbp, 5kbp, 10kbp, 25kbp, 40kbp, 50kbp, 100kbp, 250kbp, 500kbp, 1Mbp and 2.5Mbp) and normalized with ICE method ^59^. Replicate similarity was determined by HiCRep v1.12.2^60^ at 10K resolution.

For each sample, eigenvectors were determined from an ICE balanced Hi-C matrix with 40kb resolution using cooltools v0.3.2 and first principal components were used to determine A/B compartments with GC% of genome region as reference track to determine the sign. Hierarchical TADs were determined by directional index using software HiTAD v.0.4.2^61^ on bi-replicated 10kb binned matrix. Finally, for each cell type, significant intra-chromosomal interaction loops were determined under multiple resolutions (1kb, 2kb and 4kb) using the Hi-C loop callers mustache v1.0.1 (q-value < 0.1)^12^ and Fit-Hi-C2^13^ v2.0.7 (FDR < 1e-6) on merged replicates matrix. The called loops were merged between both callers to represent cell type loops at each resolution. The consensus chromatin loops within resolution were identified by combining all three cell types. These sets of loops were used as consensus for quantitative differential analysis explained below. The final consensus interaction loops for visualization (**Supplemental Figure 3E**) were collected by merging loops from all the resolutions with preference to keep the highest resolution.

Quantitative loop differential analysis across cell types was performed on fast lasso normalized interaction frequency (IF) by multiCompareHiC v1.8.0^19^ for each chromosome at resolution 1Kb, 2kb and 4kb independently. The contacts with zero interaction frequency (IF) among more than 80% of the samples and average IF less than 5 were excluded from differential analysis. The QLF test based on a generalized linear model was performed in cell type-pairwise comparisons, and p-values were corrected with FDR. The final differential loops were identified by overlapping differential IF contacts with consensus interaction loops.

### Cell-specificity score

We derived the cell type specificity weight of scRNA-seq and scATAC-seq data using CELLEX (CELL-type EXpression-specificity)^22^ on UMI counts for each gene and consensus pseudo-bulk open chromatin regions across acinar, alpha and beta cells. A gene or open chromatin region with a specificity estimate of 0 would indicate no expression in that cell type while a specificity estimate of 1 would indicate exclusive cell type-specific expression in that given cell type. The loop interaction frequency specificity was estimated by z-score across all Hi-C samples and further averaged among replicates within the same cell type. Thus, a loop with a negative z-score would indicate that it is less likely to be formed, while a positive z-score indicates loops that are more likely to formed in the given cell type.

### Transcription factor motif enrichment and cell-specific transcription factor identification

ChromVAR v1.5.0^31^ was applied to scATAC-seq to estimate transcription factor motif enrichment across pancreatic cell types. First, we constructed consensus open chromatin regions by merging pseudo-bulk open chromatin regions called for each cell type and built a cell sparse binary matrix on consensus open chromatin regions as the input for chromVAR. GC bias was corrected on BSgenome.Hsapiens.UCSC.hg19 using the addGCBias function, and the cells were filtered with a minimum fragment count of 1,500 per library and a minimum fraction of samples within peaks of 0.15. Next, we calculated the bias-corrected deviation z-scores for 746 transcription factor motifs from the nonredundant JASPAR 2020 CORE vertebrate set^32^. Motifs with differential accessibility among acinar, alpha and beta cells were identified for a given cell type compared to the other two cell types with the differentialDeviations function using a one-side t-test. Cell-specific TF motifs were determined by applying an FDR < 0.01 to the target cell type while FDR > 0.01 to the control cell types. 438 cell-specific TFs were identified by further filtering with gene expression > 0 from scRNA-seq in at least one cell type among acinar, alpha, and beta cells. To determine cell-specific TF-regulated genes, we mapped TF binding sites to the differentially accessible regions that either contact gene promoters through Hi-C loops or reside at gene promoters (-1500bp ∼ 500bp around TSS) in at least one cell type among acinar, alpha and beta cells. We performed Prerank gene set enrichment analysis with the R package fgsea^62^ using the gene expression log2 fold change the between target cell type and the other two cell types as rank.

### Bulk RNA-seq generation and analysis

Bulk RNA-seq libraries were constructed as described ^11^ and sequenced on an Illumina Novaseq 6000 instrument. The pair-end fastq files were mapped to the genome assembly hg19 by STAR (v2.6.0c) independently for each replicate. GencodeV19 annotation was used for gene feature annotation and the raw read count for gene feature was calculated by htseq-count (v0.6.1) with parameter settings -f bam -r pos -s yes -t exon -m union. The gene features localized on chrM or annotated as rRNAs, small coding RNA, or pseudo genes were removed from the final sample-by-gene read count matrix. Bulk RNA-seq data were compared to scRNA-seq to perform cell deconvolution on each donor and each cell type independently. The function music_prop from R package MuSiC v0.1.1^63^ was used to estimate the proportion of cell types from scRNA-seq in bulk RNA-seq on sorted cells. The set of selected marker features that was used for the deconvolution can be found in Supplemental Figure 2B.

### Variant to gene mapping by Hi-C and single cell data

Sentinel SNPs were obtained from the original GWAS publications^35, 37–40^ (**Supplemental Table 9**). Proxies within high LD (r2 >=0.8) with sentinel SNPs were determined using the LDlink^64^ API R package LDlinkR with the European population genotype from the 1000 Genome Project (1000G) phase 3 as reference. The proxies were filtered by overlapping with the open chromatin regions from our pseudo-bulk scATAC-seq and annotated to genes either by contacting with gene promoters through Hi-C loops or by residing at gene promoters (-1,500bp to 500bp around TSS) in the corresponding cell type (**Supplemental Figure 6**).

Variant to gene mapping was also performed using the Activity-by-Contact (ABC) model^41^ to link SNP-containing putative enhancers to target genes with the cell type-specific Hi-C matrix. ATAC-seq signals were obtained by pooling fragment mapping of scATAC-seq in acinar, alpha and beta cells, respectively, while the H3K27ac ChIP-seq signals were derived from publicly available resources for acinar (GEO:GSE79468)^16^, alpha and beta cells (GEO:GSE140500)^65^, and pre-processed using the Encode ChIP-seq pipeline (https://github.com/ENCODE-DCC/chip-seq-pipeline2). The candidate enhancer regions were identified and quantified by mapping scATAC-seq and H3K27ac ChIP-seq signals to pseudo bulk open chromatin regions merged from acinar, alpha and beta cells, and quantile normalized against the K562 cell line epigenetic signal provided by the software. Protein-coding and long non-coding RNA gene promoters were defined by Gencode V19 annotation with ∼500bp around TSS. The ABC scores were calculated for each candidate region-to-transcript pair by integrating cell type-specific Hi-C contact frequencies at 4kb resolution from our acinar, alpha and beta cell data. Any putative enhancer-to-transcript pair with an ABC score above 0.02 was identified as a significant interaction and used to overlap with T2D risk proxies as identified above.

Disruptive effect of T2D putative causal variants on JASPAR 2020 transcription factor binding sites was estimated using motifbreakR^66^ with relative entropy algorithm (method = “ic”. The results were filtered with PWM match p-value smaller than 1e-3 and TF gene expression > 0 in corresponding cell types.

### Partitioned heritability LD score regression enrichment analysis

Partitioned heritability LD Score Regression (v1.0.0)^34^ was used to identify heritability enrichment with GWAS summary statistics and open chromatin regions annotated to genes. The baseline analysis was performed using LDSCORE data (https://data.broadinstitute.org/alkesgroup/LDSCORE) with LD scores, regression weights, and allele frequencies from the 1000G phase 1 data. The summary statistics were obtained from GWAS publications^33, 35, 37, 39, 40^ and harmonized. The Partitioned LD score regression annotations were generated using the coordinates of open chromatin regions which either contact gene promoters through Hi-C loops or reside at gene promoters for each cell type. Finally, partitioned LD scores were compared to baseline LD scores to measure enrichment fold change and enrichment p-values, which were adjusted with FDR across all comparisons.

### Multi-trait colocalization analysis

Sentinel signals in T2D GWAS were obtained from putative causal variant to gene pairs shared between T2D and other relevant diabetes traits and used as input SNPs to identify multi-trait colocalized association signals. Summary statistics on different traits were harmonized to the 1000 genome reference data set and the missing variants were imputed using scripts from https://github.com/hakyimlab/summary-gwas-imputation. The effect size and standard deviation for imputed variants were calculated from estimated z-scores, reference allele frequencies and effective sample sizes^67^. The colocalization analysis was performed for 1Mbp regions around each input sentinel signal using HyPrColoc^68^, and the posterior probability of colocalization at putative causal variants identified by variant-to-gene mapping was approximated by setting snpscores as TRUE in the hyprcoloc function.

## Supplemental Figures

**Supplemental Figure 1. Sorting of the major pancreatic cell populations.** Representative FACS plots showing the gates use to select acinar, alpha and beta cell populations.

**Supplemental Figure 2. Pancreatic islet cell transcriptomic profile defined using scRNA-seq. A**, Clustering of gene expression profile from 5,288 pancreatic cells from non-diabetic organ donors identified 9 distinct clusters on UMAP. Endo: endothelial; eps: epsilon (ghrelin producing); mes: mesenchymal; PP: (pancreatic polypeptide producing). **B,** the expression of marker genes in single cells and donor composition within cell type clusters. **C,** Pearson correlation of gene expression between donors. **D,** Pearson correlation of gene expression between cell types. **E,** Estimate of cell composition in sorted pancreatic cell subsets through integration of scRNA-seq. The cell type composition from bulk RNA-seq data on sorted pancreatic cell subsets were characterized using a list of gene markers obtained from panel B.

**Supplemental Figure 3. Pancreatic islet cell chromatin accessibility profile defined using scATAC-seq. A,** Clustering of accessible chromatin profile from 12,473 pancreatic cells non-diabetic organ donors identified 9 distinct clusters on UMAP. **B**, Promoter accessibility of selected marker genes in single cells and donor composition within cell type clusters. Promoters were defined as the window of -1,500 to +500bp around TSS in Gencode V19 annotation. **C**, scATAC-seq and bulk ATAC-seq accessibility signal around selected marker genes: CPA1 (acinar), GCG (alpha) and IGF2 (beta). Fragments were aggregated for each cell type in scATAC-seq. The coverage for both scATAC-seq and bulk ATAC-seq was normalized using the RPGC method from deeptools. **D**, Pearson correlation of OCR accessibility between scATAC-seq and bulk ATAC-seq. OCR accessibility were calculated as FPKM by mapping either cell type aggregated fragments from scATAC-seq or fragments of bulk ATAC-seq to the consensus pseudo-bulk peak reference. **E**, Spearman correlation between t-statistics of cluster-specific genes based on promoter accessibility (scATAC-seq) and gene expression (scRNA-seq).

**Supplemental Figure 4. 3D chromatin architecture of pancreatic cell subsets defined by Hi-C assay. A**, Stratum-adjusted correlation coefficient (SCC) between Hi-C samples to measure sample reproducibility and interrelationships of cell lineages. **B**, the number of Topologically Associating Domains (TADs) identified in each cell type. **C**, the distribution of TAD size in each cell type. **D**, Aggregate peak analysis (APA) plots of chromatin loops identified by the loop callers Mustache and Fit-Hi-C2. The APA scores were calculated with respect to the enrichment of the center pixel at 4kb resolution and labelled at the top left corner. **E**, Venn diagram of consensus loop calls across cell types. The consensus loops were collected by merging loop calls from 1kb, 2kb and 4kb resolutions with preference to keep the highest resolution data. **F**, The number of loops annotated to open chromatin regions (OCR) and genes. OCR (green): both anchors overlapped OCRs; Gene (teal): both anchors overlapped gene promoters (-1,500 to +500bp around the TSS). Gene-OCR (purple): one anchor overlapped an OCR and the other anchor overlapped a gene promoter. **G,** Z-score distribution of co-accessible promoters and distal OCRs detected by Hi-C chromatin loops. P-values were calculated by two-sided Wilcoxon rank sum test. **H,** The number of loop anchors overlapped with chromHMM chromatin features. chromHMM chromatin states for acinar and endocrine cells were previously defined in Arda *et al.* 2018. **I,** Enrichment of chromatin loops at open chromatin regions and regulatory chromatin features defined by chromHMM. The enrichment fold change was obtained by comparing called loops to distance-matched random chromatin contacts. The p-value was calculated by a 100-fold permutation test.

**Supplemental Figure 5. Transcription factor (TF) modules and TF-gene networks for beta- and acinar-cell specific TFs. A and B**, Association index analysis revealed acinar- (**A**) and beta-cell (**B**) specific TF modules. TF sets are defined as cell-specific TFs with top 30 significant GSEA enrichment (**Supplemental Table 7**). Matrices show the Jaccard similarity between TF pairs based on occurrence of co-regulated downstream genes. **F**, TF-gene network for the top 30 beta-cell specific TF sets. The nodes are either TF (green triangle) or downstream genes (red circle) from the leading edge in **Supplemental Table 7**. The edges represent how TFs and downstream genes are connected: TF motif at only distal cis-regulatory elements (red), TF motif at only gene promoters (green), and TF motif at both gene promoter and connected cis-regulatory element (blue).

**Supplemental Figure 6. Illustration of variant-to-gene mapping**. The candidate variants were first defined by proxy SNPs within high LD (R2 > 0.8) with sentinel signals from GWAS, and then filtered to proxy SNPs located in open chromatin by peaks called from ATAC-seq. The open proxy SNPs were mapped to candidate genes with two approaches. (1) Distal genes were linked to open SNP-containing peaks with physical interaction evidence from Hi-C loop calls for each cell type (left diagram). (2) Proximal genes whose promoter regions (-1,500bp ∼ +500bp of TSS) fall within open SNP-containing peaks.

**Supplemental Figure 7. Comparison of T2D variant-to-gene mappings using the ABC model and chromatin loop binary calls. A,** Venn diagram of gene to distal-element pairs. The gene-distal-element pairs were identified independently in each cell type using the ABC model or our Hi-C chromatin loops (see **Methods**), and a union list was obtained. The number of shared gene-distal-element pair and the percentage of intersect over the union between the two approaches are labeled. **B**, Venn diagram of variant-to-gene mapping results in T2D. The putative causal-variant to gene pairs were defined by overlapping gene-distal-element maps with open proxies within high LD (R2>2) with T2D sentinel signals. The number of candidate genes, causal-variant-containing OCRs, candidate causal variants, causal-variant-gene pairs, corresponding sentinel signals and sentinel-signal-gene pairs were summarized in Venn diagram. **C**, Specificity and sensitivity of variant-to-gene mapping approaches using chromatin maps from pancreatic cells to detect positive control genes previously defined for T2D. The approaches “ABC” and “binary” were detailed in Methods. The “open promoter” approach defined candidate genes as the proximal genes whose promoter regions (-1,500bp to +500bp of TSS) overlapped with proxies located in open chromatin regions (**Supplemental Figure 6** right branch), while the “open nearest genes” were defined by proximal genes closest to open proxies regardless of distance.

**Supplemental Figure 8.** Examples of likely disruption of transcription factor DNA binding motifs by T2D relevant SNPs. A. Variants at the TH/INS/TRPM5 locus that are in contact loops with the INS or TRPM5 promoter likely disrupt binding of the TFs indicated. B. Disruption of a putative NEUROD2 binding site by rs4234731 in the WSF1 gene.

**Supplemental Figure 9. Comparison of T2D variant-to-gene mapping with other diabetes relevant traits. A**, ‘Upset’ plot of putative casual-variant-gene pairs across diabetes-related traits. Examined traits include type 2 diabetes (T2D), fasting glucose (FG), Hemoglobin A1C (HbA1c), fasting insulin (FI), 2h glucose (2hGlu) and type 1 diabetes (T1D). Each row corresponds to the set of putative casual-variant to gene pairs in each trait. Each column corresponds to one segment of a Venn diagram obtained by intersecting the traits indicated as black circles. All segments that intersect with T2D are highlighted in red. The number of putative casual-variant to gene pairs in each set and intersecting segment are summarized in the bar graph on top. **B**, Stacked association plots of T2D with FG and HbA1c at T2D sentinel signal rs10228066. Variant-to-gene mapping implicates rs10228796 as the candidate causal variant and the linked rs10228796 to the candidate effector gene *DGKB* with beta cell-specific chromatin loops (**Figure 5E**, **Supplemental Table 10**). **C**, HyPrColoc identified rs10228796 as a candidate causal variant explaining the shared association signal among T2D, FG and HbA1c. The posterior probability of colocalization between the traits was 0.819 and rs10228796 explained 33.6% of their colocalization

## Supplemental Tables

Supplemental Table 1. Donor information for experimental assays.

Supplemental Table 2. Hi-C library hicup pre-processing summary

Supplemental Table 3. Hi-C chromatin loop call summary

Supplemental Table 4. GO enrichment of genes under acinar- and endocrine-specific A compartments.

Supplemental Table 5. Clustering of Cell-type differential loops (CTDLs).

Supplemental Table 6. Cell-specific TF motif chromVAR enrichment

Supplemental Table 7. GSEA enrichment of downstream genes regulated by cell-specific TFs

Supplemental Table 8. GO enrichment of genes implicated by V2G mapping in T2D

Supplemental Table 9. GWAS sentinel signals of T2D, T1D and glycemic traits.

Supplemental Table 10. Variant-to-gene mapping result in T2D

Supplemental Table 11. Predicted disruption of TF binding motifs by putative causal variants for T2D

Supplemental Table 12. Implicated effector genes from our variant to gene mapping approach, and the corresponding observed cell type(s).

Supplemental Table 13. Variant-to-gene mapping result in T1D and glycemic traits

Supplemental Table 14. Multiple trait colocalization between T2D and relevant traits.

