## Supplementary figures and images for "The three-dimensional chromatin structure of the major human pancreatic cell types reveals lineage-specific regulatory architecture of T2D risk"

### Supplemental Figure 1

#### Supplemental Figure 1

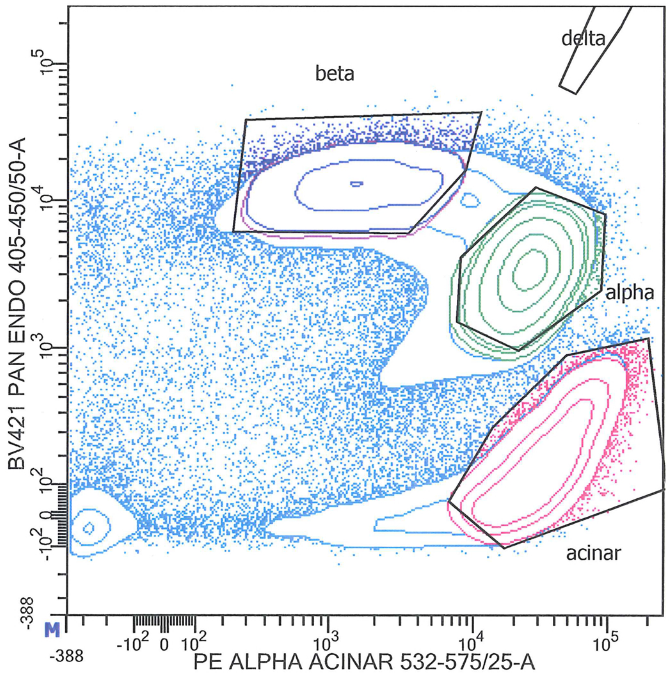


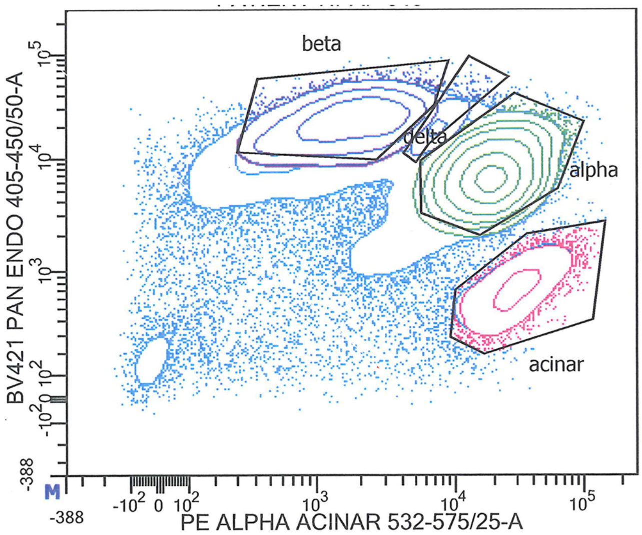

### Supplemental Figure 2

**Supplemental Figure 2.**

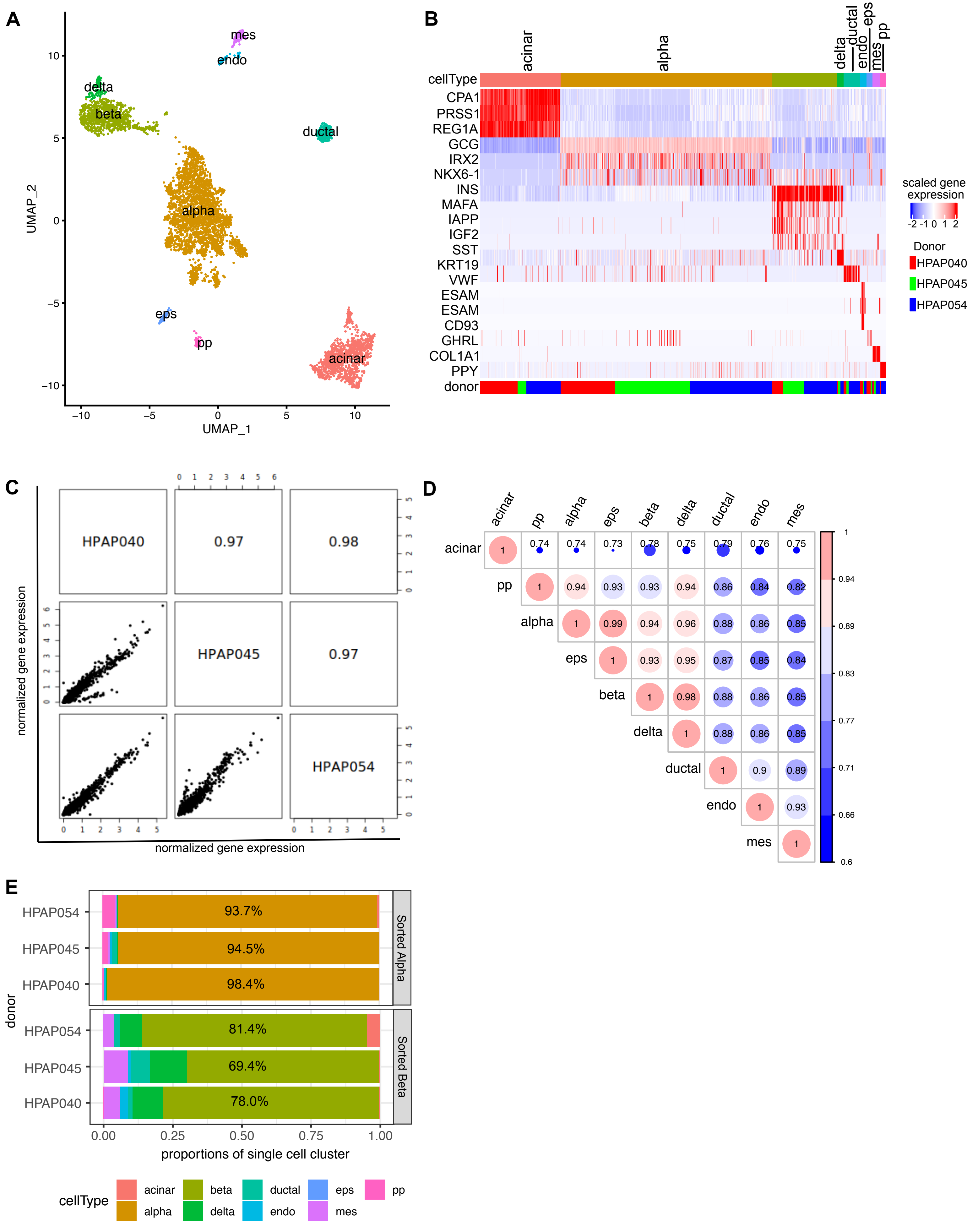

### Supplemental Figure 3

##### Supplemental figure 3

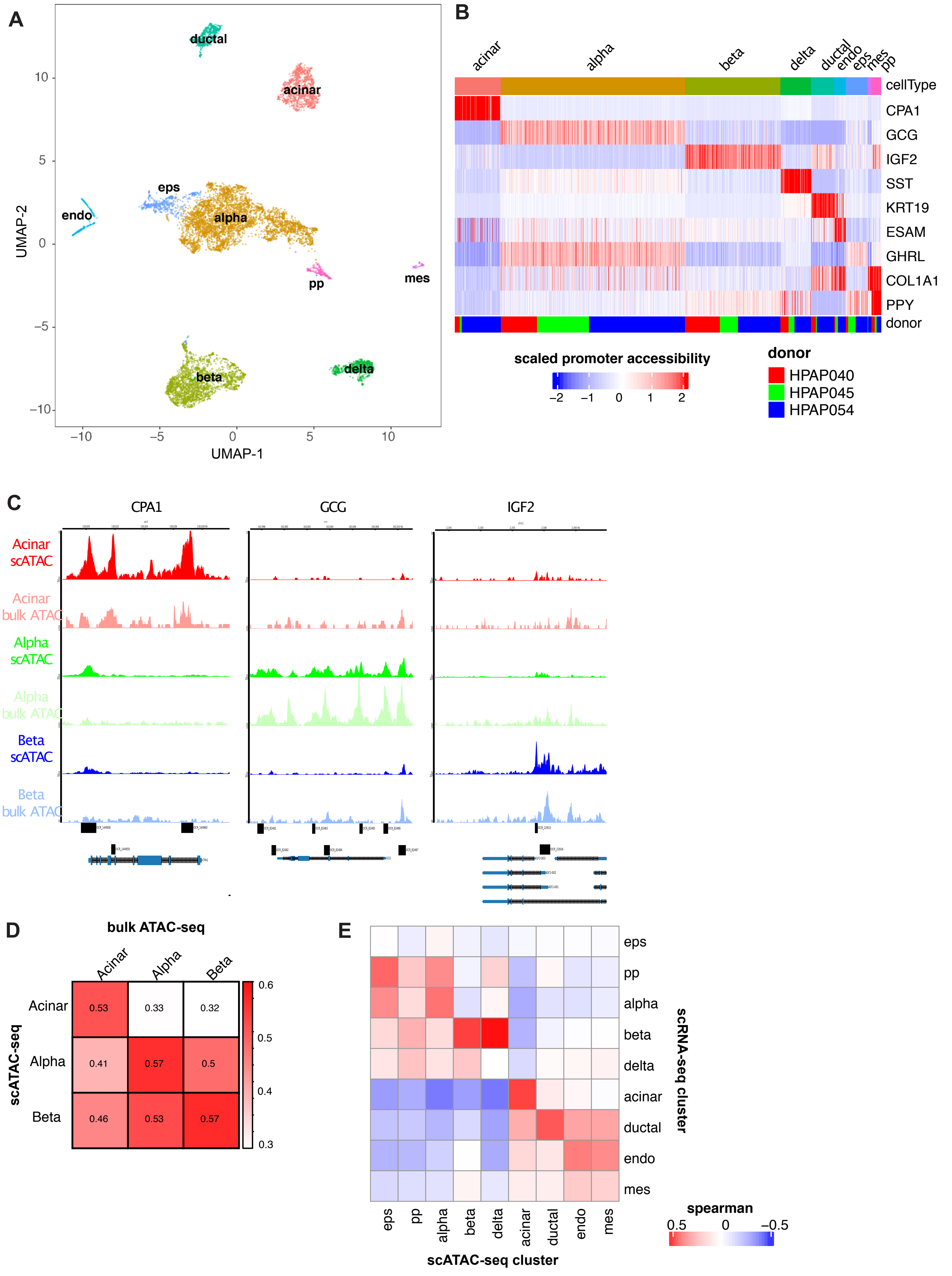

### Supplemental Figure 4

#### Supplemental Figure 4

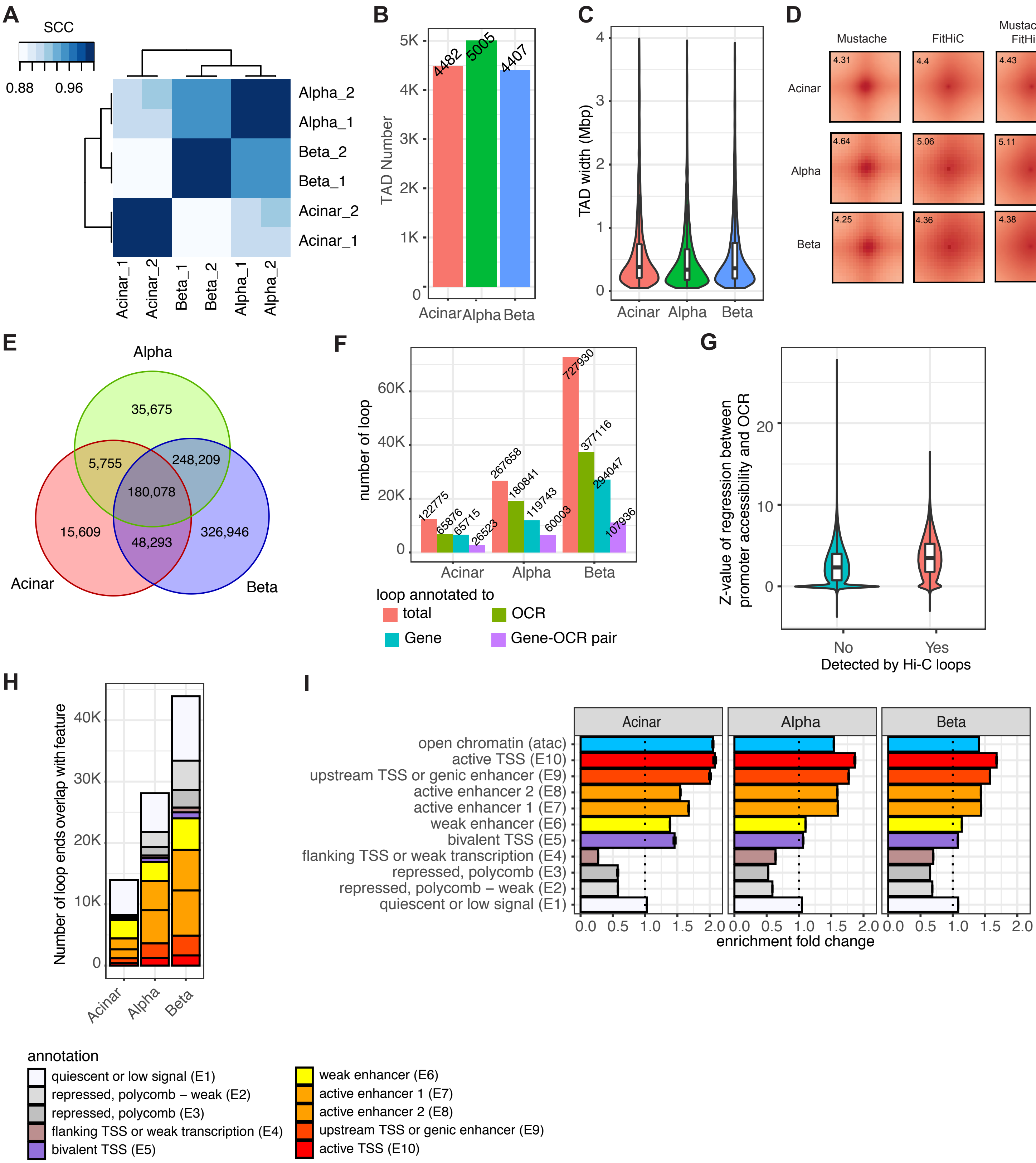

### Supplemental Figure 5

##### Supplemental Figure 5

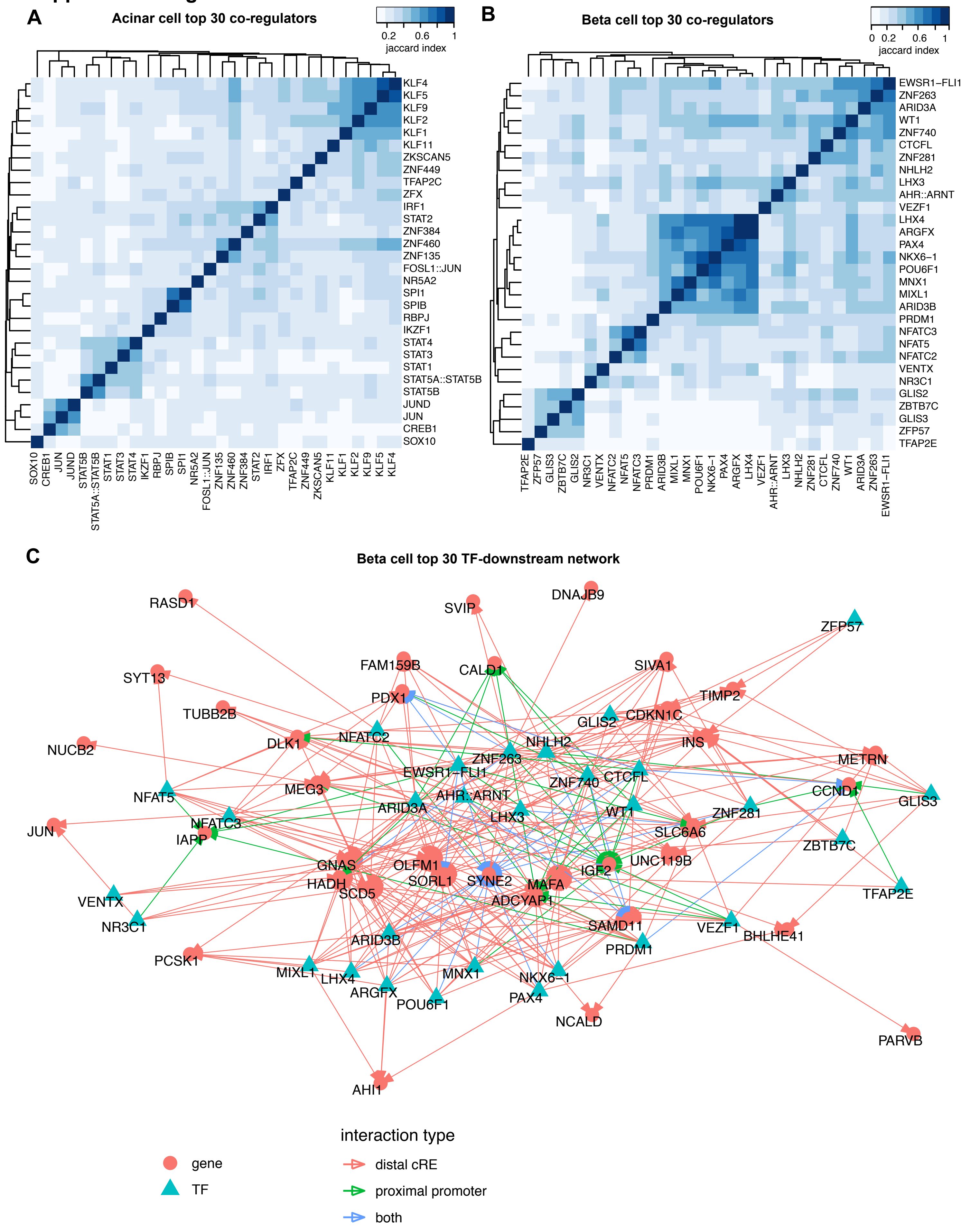

### Supplemental Figure 6

##### Supplemental Figure 6

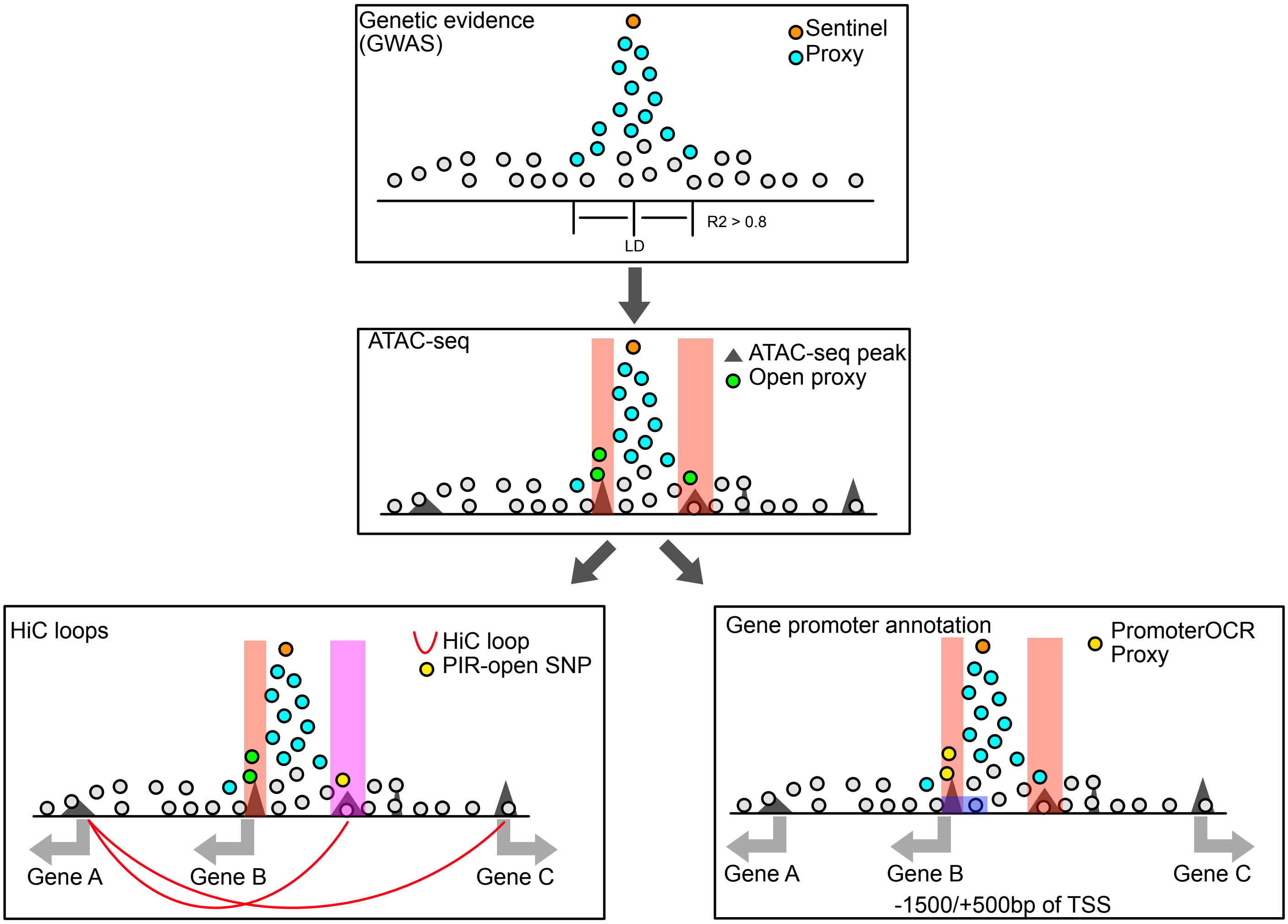

### Supplemental Figure 7

##### Supplemental Figure 7

**A.**

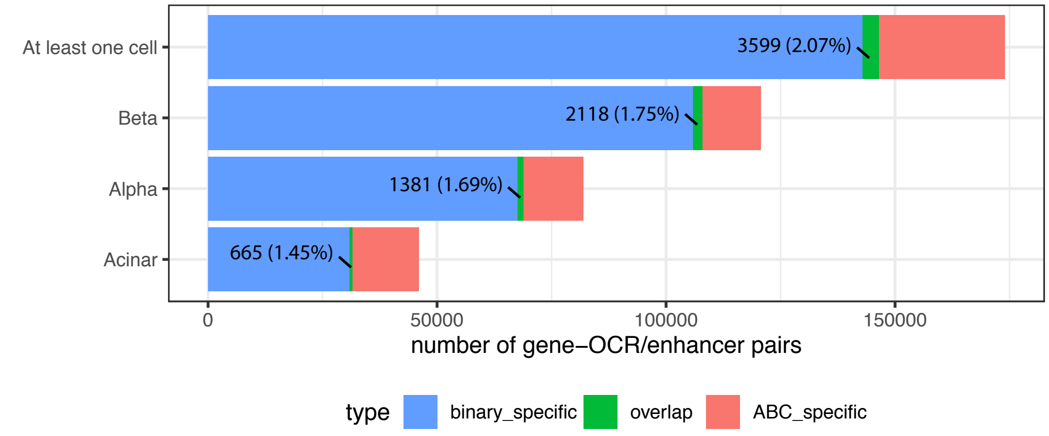


**B.**

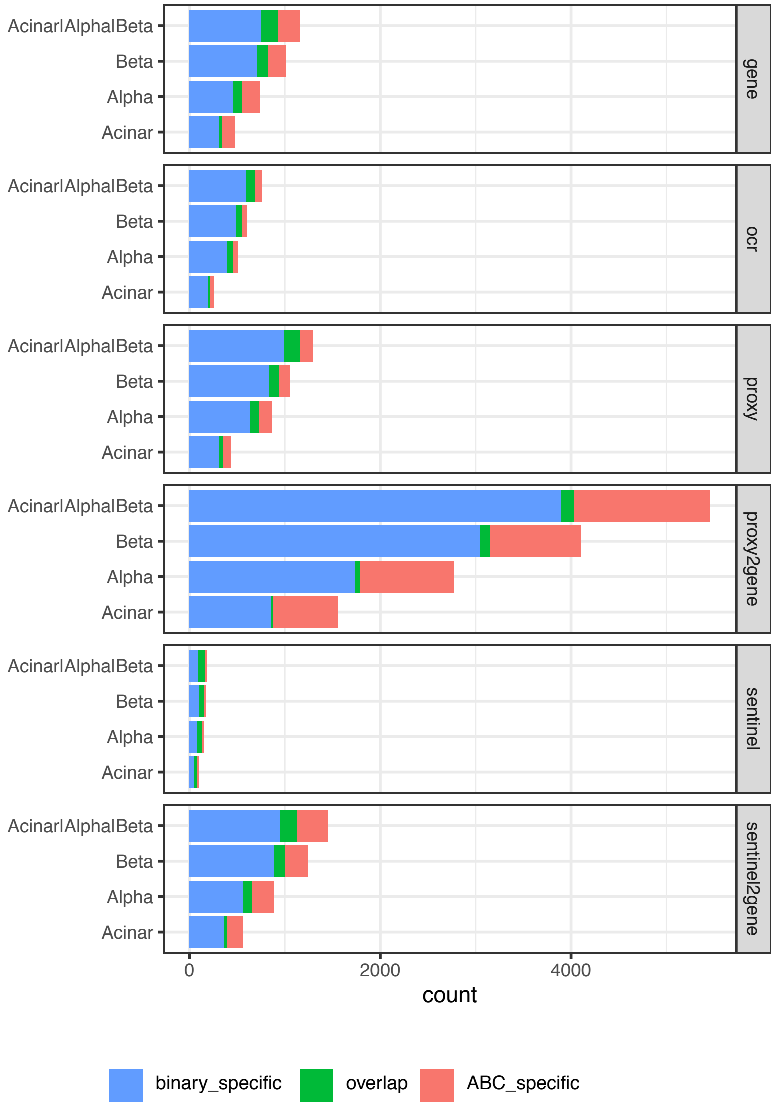

### Supplemental Figure 8

#### Supplemental Figure 8

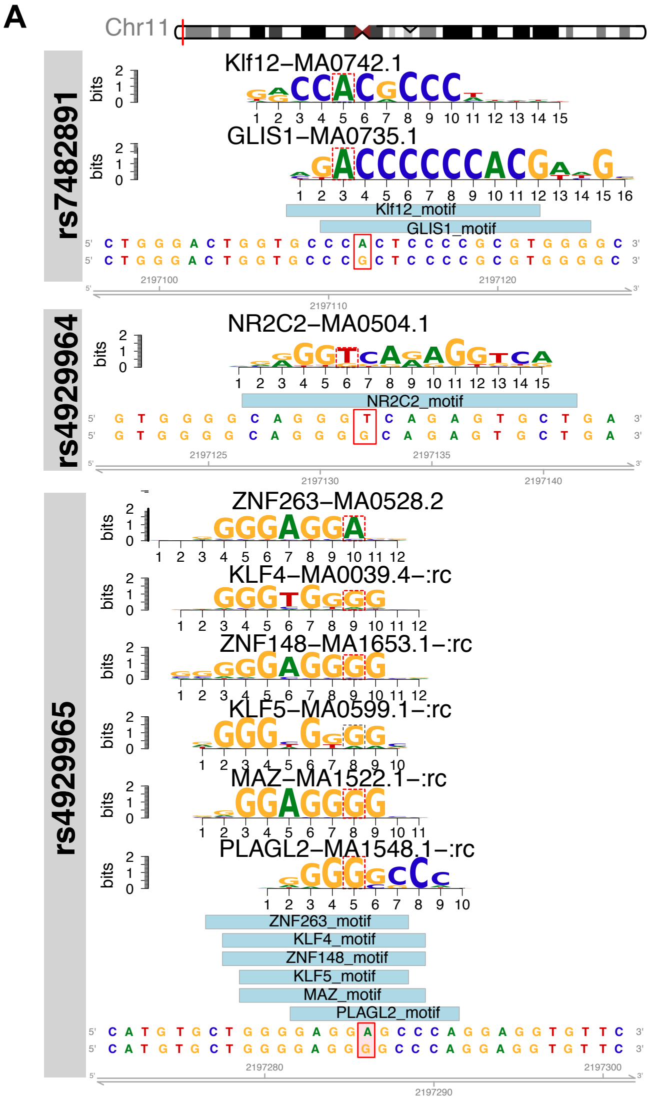


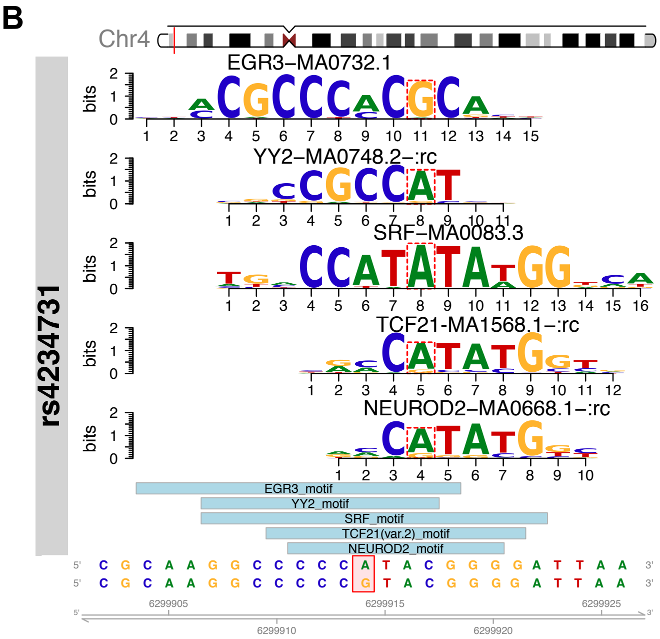

### Supplemental Figure 9

#### Supplemental Figure 9

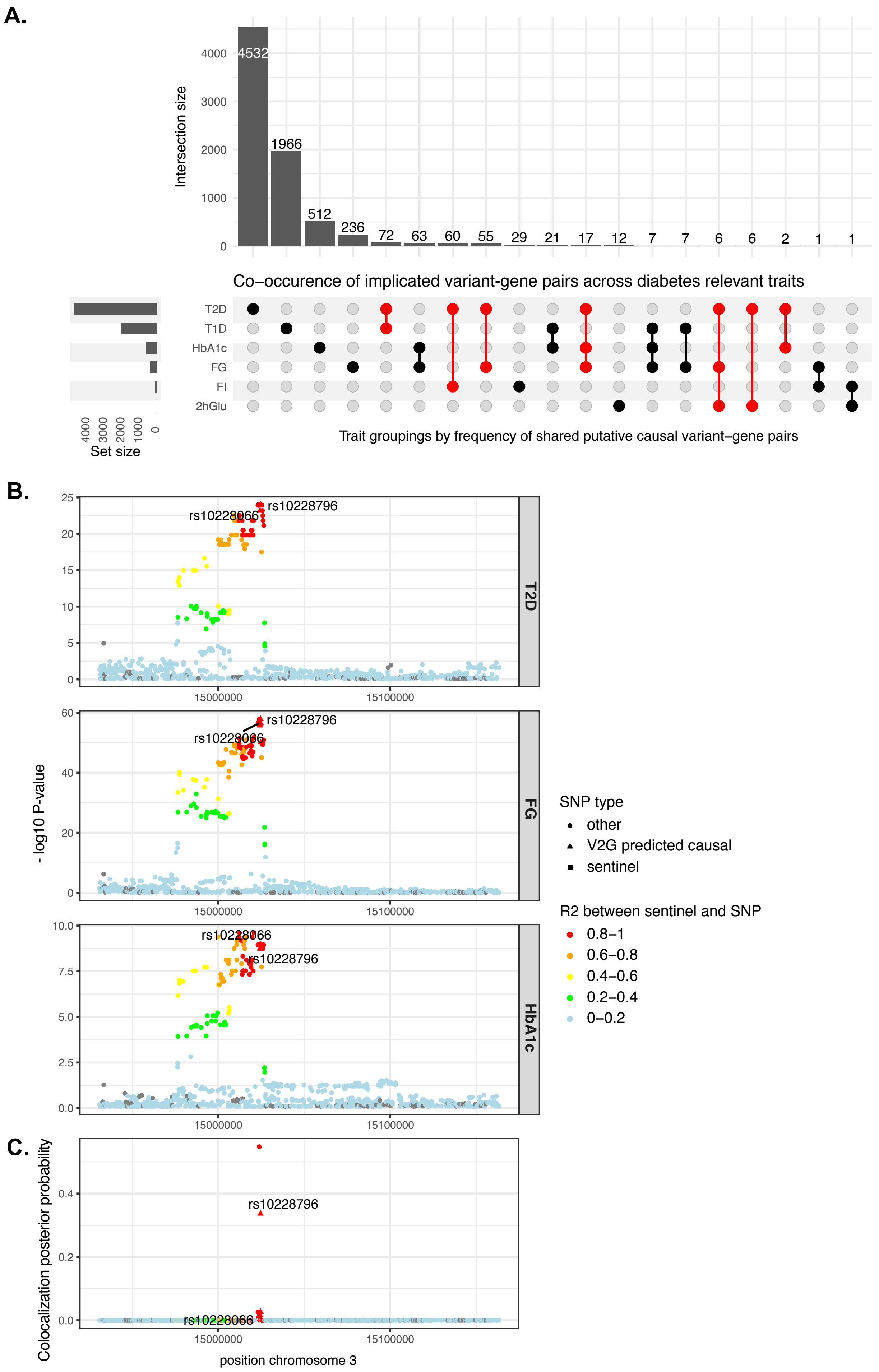
